# SON and SRRM2 form nuclear speckles in human cells

**DOI:** 10.1101/2020.06.19.160762

**Authors:** İbrahim Avşar Ilık, Michal Malszycki, Anna Katharina Lübke, Claudia Schade, David Meierhofer, Tuğçe Aktaş

## Abstract

The nucleus of higher eukaryotes is a highly compartmentalized and dynamic organelle consisting of several biomolecular condensates that regulate gene expression at multiple levels (1, 2). First reported more than 100 years ago by Ramon y Cajal, nuclear speckles (NS) are among the most prominent of such condensates (3). Despite their prevalence, research on the function of NS is virtually restricted to colocalization analyses, since an organizing core, without which NS cannot form, remains unidentified (4, 5). The monoclonal antibody SC35, which was raised against a spliceosomal extract, is a frequently used reagent to mark NS since its debut in 1990 (6). Unexpectedly, we found that this antibody has been misidentified and the main target of SC35 mAb is SRRM2, a large (∼300 kDa), spliceosome-associated (7) protein with prominent intrinsically disordered regions (IDRs) that sharply localizes to NS (8). Here we show that, the elusive core of NS is formed by SON and SRRM2, since depletion of SON leads only to a partial disassembly of NS, while combined depletion of SON together with SRRM2, but not other NS associated factors, or depletion of SON in a cell line where IDRs of SRRM2 are genetically deleted, leads to a near-complete dissolution of NS. This work, therefore, paves the way to study the role of NS under diverse physiological and stress conditions.

## Introduction

Nuclear speckles (NS) are membraneless nuclear bodies in the interchromatin-space of the nucleus that contain high concentrations of RNA-processing and some transcription factors but are devoid of DNA (3). Under normal conditions, they appear as irregularly shaped, dynamic structures that show hallmarks of phase-separated condensates, such as fusion and deformation under pressure in living cells (5, 9). Despite their prevalence, the function of NS remains largely unknown, although they have been proposed to act as reservoirs for splicing factors, and association with NS have been shown to correlate with enhanced transcription and RNA processing (4, 5). NS have been shown to be involved in replication of herpes simplex virus (10), processing and trafficking of Influenza A virus mRNA (11), detaining repetitive RNA originating from the transcription of repeat expanded loci that trigger Huntington’s disease, spinocerebellar ataxia and dentatorubral–pallidoluysian atrophy (12), but also repetitive RNA from artificial constructs that produce RNA capable of phase-separation in vitro (13). Studying the role of NS involves visualizing them with a fluorescently tagged factor that localizes to NS, or use the of antibodies that show specific staining of NS. Similar to nucleoli and other membraneless bodies of the nucleus, NS disassemble during early stages of mitosis, and assemble back following telophase (4). Several protein kinases are thought to be involved in this process, such as DYRK3, chemical inhibition of which leads to aberrant phase-separation (14). Overexpression of DYRK3, or CLK1 on the other hand leads to dissolution of NS in interphase cells, underscoring the importance of phosphorylation in NS integrity (14, 15). Unlike several other biomolecular condensates, a specific core necessary for NS formation has not yet been identified, and it has been hypothesized that stochastic self-assembly of NS-associated factors could lead to the formation of NS (3, 16, 17). One of the most frequently used reagents to locate NS is the monoclonal antibody SC35, which was raised against biochemically purified spliceosomes (6), and reported to be an antibody against SRSF2 (18). Testament to the importance of this antibody, NS are also referred to as ‘SC35 domains’. Although, the name SC35 and SRSF2 are used synonymously and to annotate orthologues of SRSF2 not only in mammalian species but also in species such as *D. melanogaster* and *A. thaliana*, mAb SC35 is reported to cross-react with SRSF1, and potentially with other SR-proteins as well (19). Furthermore, fluorescently tagged SRSF2 shows staining patterns incompatible with mAb SC35 stainings under identical experimental conditions (20–23). Intrigued by these inconsistencies, we undertook a systemic re-characterization of the mAb SC35 and its cellular targets.

## Results

### IP-MS reveals endogenous targets of mAb SC35

In order to characterize the cellular targets of the SC35 mAb, we carried out an Immunoprecipitation Mass-Spectrometry (IP-MS) experiment. Whole-cell extracts prepared from HAP1 cells were used to immunoprecipitate endogenous targets SC35 mAb, with a matched IgG mAb serving as a control. The immunoprecipitated proteins were then analyzed by mass-spectrometry (see Methods for details). In total, we identified 432 proteins that were significantly enriched in the SC35 purifications compared to controls (p<0.05, at least 2 peptides detected in each biological replicate). Surprisingly, the most enriched protein in the dataset, both in terms of unique peptides detected, total intensities and MS/MS spectra analyzed, is neither SRSF2 nor one of the canonical SR-proteins (24), but a high-molecular weight RNA-binding protein called SRRM2 (Fig. 1A, Supplementary Fig. 1A), an NS-associated protein with multiple RS-repeats (25). Analysis of the top 108 targets, corresponding to the third quartile, using the STRING database (26) shows a clear enrichment for the spliceosome and nuclear speckles (Supplementary Fig. 1B), validating the experimental approach. We were also able to detect all SR-proteins in our dataset, however their scores are dwarfed by that of SRRM2’s (Supplementary Fig. 1C). Thus, contrary to initial expectations, the IP-MS results thus strongly suggest that SC35 mAb primarily recognizes SRRM2, at least in the context of an immunoprecipitation experiment.

**Fig. 1.**
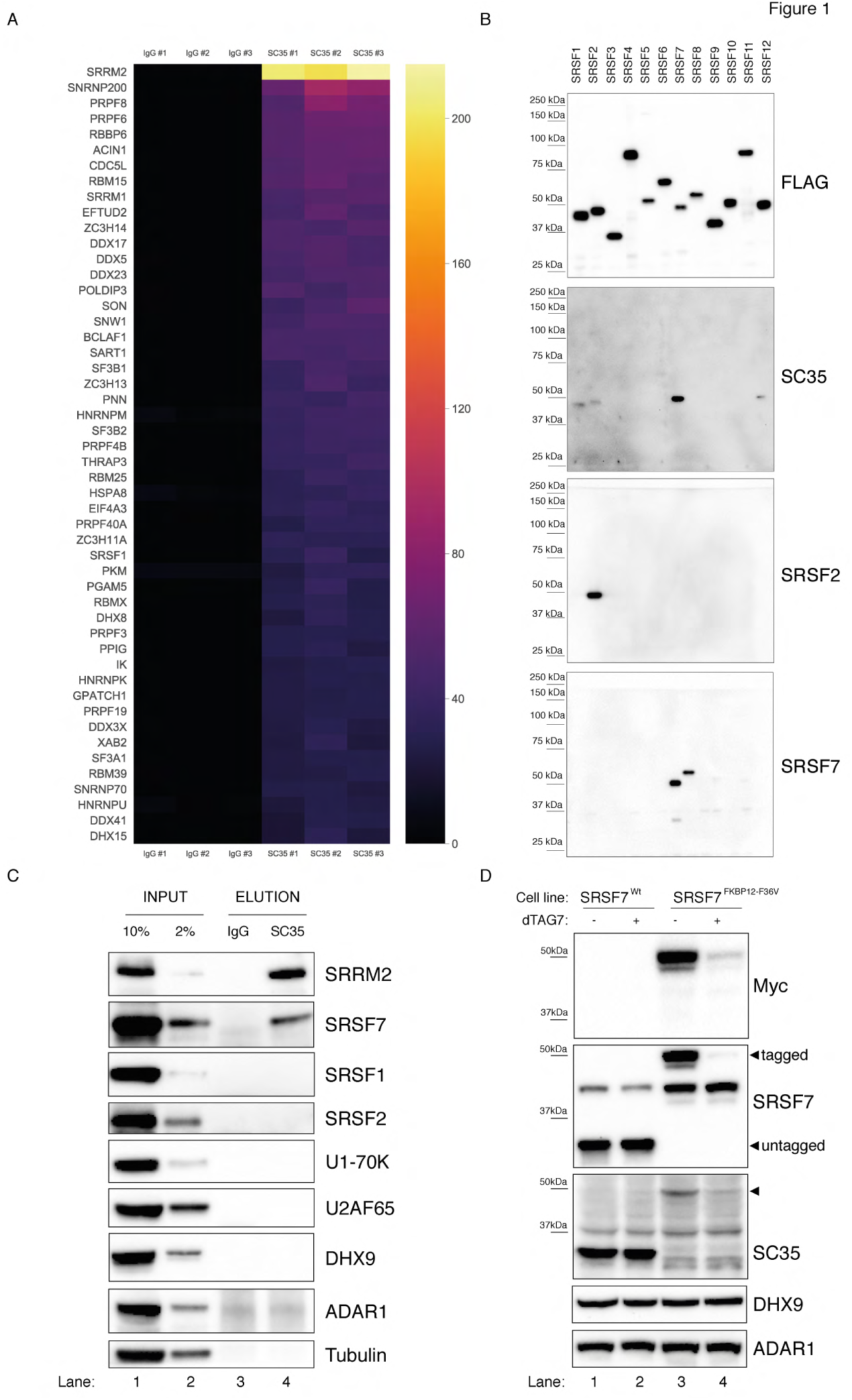
SC35 mAb immunoprecipitation followed by MS identifies SRRM2 as the top hit. **(A)** The Top50 hits identified by the MS are depicted on a heatmap showing the number of unique peptides detected for each protein. Also see Supplementary Fig. 1A for an intensity vs MS/MS spectra plot and Supplementary Table 2 for raw MaxQuant results **(B)** Streptavidin pull-down of biotin tagged ectopically expressed SRSF proteins 1 to 12 are blotted with FLAG antibody in order to show the amounts of loaded proteins on PAGE. Western blot using mAb SC35 detects SRSF7 with highest sensitivity in comparison to all other SRSF proteins, but also weakly reacts with SRSF1, 2 and Specific antibodies against SRSF2 and SRSF7 are used to validate the authenticity of the purified proteins from stable cell lines in corresponding lanes and blots. **(C)** SC35-IP performed on lysates from wild-type HEK293 cells identifies SRRM2 as the most enriched protein with a weaker enrichment for SRSF7 but no enrichment for SRSF1 or SRSF2 using western blotting (compare Lane 4 across blots). **(D)** Homozygous knock-in of the 2xMyc-FKBP12^F36V^ tag into *SRSF7* gene locus shifts the SC35 band from 35kDa to 50kDa (compare Lanes 1 and 3 on the SC35 blot) and upon induction of degradation with dTAG7 the shifted band is lost (compare Lanes 3 and 4 in Myc, SRSF7 and SC35 blots). This blot validates that the 35 kDa band identified by mAb SC35 blots corresponds to SRSF7.

### SRSF7 is the 35 kDa protein recognised by mAb SC35

Before exploring SRRM2 as a potential mAb SC35 target protein, we decided to first take an unbiased look at SR-proteins and the ability of mAb SC35 to recognize them. To this end, we cloned all 12 canonical SR-proteins in humans (24) into an expression plasmid, and created stable-cell lines expressing these proteins under the control of a doxycycline-inducible promoter (Supplementary Fig. 2). We used a biotin-acceptor peptide as a tag, and carried out stringent purifications using streptavidin beads to exclude non-specific co-purification of unrelated SR-proteins, and examined the eluates using immunoblotting. Surprisingly, our results show that the main target of mAb SC35 on these immunoblots is SRSF7 (Fig. 1B), which runs at approximately ∼35kDa on polyacrylamide gels similar to SRSF2. To exclude any artefacts that could originate from the use of tagged proteins, we used whole-cell extracts from HEK293 cells and immunoprecipitated targets of mAb SC35 and analyzed the eluates via immunoblotting. Consistent with the results of the IP-MS experiment, and tagged-SRSF1-12 purifications, we observed a very clear enrichment for SRRM2 and SRSF7, but not for SRSF2, SRSF1 or other factors (Fig. 1C). In order to determine whether the 35 kDa band recognized by mAb SC35 in immunoblots of cellular lysates is composed of multiple proteins, in addition to SRSF7, we created a cell line where we inserted the FKBP12^F36V^ degron (27) homozygously into the C-terminus of SRSF7 in HEK293 cells. Even without any treatment, it is evident that the 35 kDa band robustly recognized by mAb SC35 in wild-type cells completely disappears in SRSF7^FKBP12^ cells (Fig. 1D, compare lanes 1 and 2 with 3 and 4), and a new band around ∼50 kDa, where the FKBP12^F36V^-tagged SRSF7 runs, emerges (Fig. 1D, arrow-head). Treatment of these cells with 1µM of dTAG7 for 6hrs lead to the depletion FKBP12^F36V^-tagged SRSF7, and to the depletion of the newly-emerged ∼50 kDa protein recognized by mAb SC35. These results strongly suggest that the 35 kDa namesake protein revealed by SC35 mAb on immunoblots is SRSF7 and any contribution to this signal from other proteins is negligible to none.

### SRRM2 is the primary target of mAb SC35 in immunoblots

Even though our results show that SC35 mAb specifically recognizes SRSF7 rather than SRSF2, both proteins have significant nucleoplasmic pools in addition to their localization to nuclear speckles (23, 28, 29) which is not easily reconciled with the immunofluorescence stainings obtained with the SC35 mAb that are virtually restricted to nuclear speckles. Intriguingly, SRRM2, which is by far the most enriched protein in our immunoprecipitations with mAb SC35, is a relatively large (∼300 kDa) protein, that readily co-purifies with spliceosomes (30, 31), shows liquid-like behaviour in cells (14) and co-localizes near-perfectly with mAb SC35-stained NS (32). In addition, SRRM2 and its yeast counterpart Cwc21/Cwf21 is located in the recent cryoEM structures of the spliceosome, where it joins the spliceosome at the B^act^ stage where it seems to support the activated conformation of PRP8’s switch-loop both in humans and yeast (7). Predating the recent cryo-EM structures by almost a decade, the yeast orthologue of SRRM2, Cwf21p, has been shown to directly interact with Prp8p (PRPF8) and Snu114p (EFTUD2) which are also among the most enriched proteins in our mAb SC35 immunoprecipitations (33)(Supplementary Fig. 1A). Furthermore, a more recent tandem-affinity purification of the protist *Trypanosoma* orthologue of SRRM2 revealed Prp8, U5-200K (SNRNP200, also known as Brr2), U5-116K (EFTUD2, also known as Snu114) and U5-40K (SNRNP40) as major interaction partners (34).

Taking into consideration the fact the mAb SC35 was raised against biochemically purified spliceosomes (6), together with the aforementioned observations in the scientific literature and our IP-MS results which identified SRRM2 as the top target, we hypothesize that mAb SC35 was most likely raised against SRRM2, and it recognizes SRRM2 in most if not all immunological assays that utilizes mAb SC35 where SRRM2 is not depleted or unintentionally omitted due to technical reasons.

In order to test the veracity of this claim, we designed a series of experiments in human cells. Since, to our knowledge, SC35 mAb has not been shown to recognize SRRM2 on immunoblots, we first created tagged and truncated SRRM2 constructs in living cells. To this end, we generated 11 cell lines that remove between 4 and 2,322 amino acids from the SRRM2 protein (full-length: 2,752 a.a., numbering from Q9UQ35-1) by inserting a TagGFP2 (referred to as GFP for simplicity) sequence followed by an SV40 polyadenylation signal into 11 positions of the *SRRM2* gene in HAP1 cells using a CRISPR/Cas9-based technique called CRISPaint (35) (Fig. 2A). The deepest truncation removes 84% of SRRM2, which includes almost all its IDRs, together with two regions enriched for serine and arginine residues, leaving behind 13 RS-dinucleotides out of a total of 173 (Supplementary Fig. 3A,B). The GFP-tagged, in vivo truncated proteins (referred to as **tr**uncations 0 to 10, shortened as tr0 - tr10, Fig. 2B) are then immunoprecipitated using GFP-trap beads and the eluates were analyzed by immunoblotting. This experiment shows that SC35 mAb indeed recognizes SRRM2 on immunoblots (Fig. 2C, lane 2). Interestingly, the signal from SC35 mAb remains relatively stable up until SRRM2^tr4^ which removes 868 a.a. from the SRRM2 C-terminus, the signal appears to be reduced in SRRM2^tr5^ which removes 1,014 a.a. and becomes completely undetectable from SRRM2^tr6^ onward (Fig. 2C and Supplementary Fig. 3C). The same blot was stripped and re-probed with a polyclonal antibody raised against the N-terminus of SRRM2, common to all truncations, which show that SRRM2 is detectable throughout, and thus indicating that the epitope(s) recognised by mAb SC35 reside between amino acids 1,360 - 1,884 of SRRM2.

**Fig. 2.**
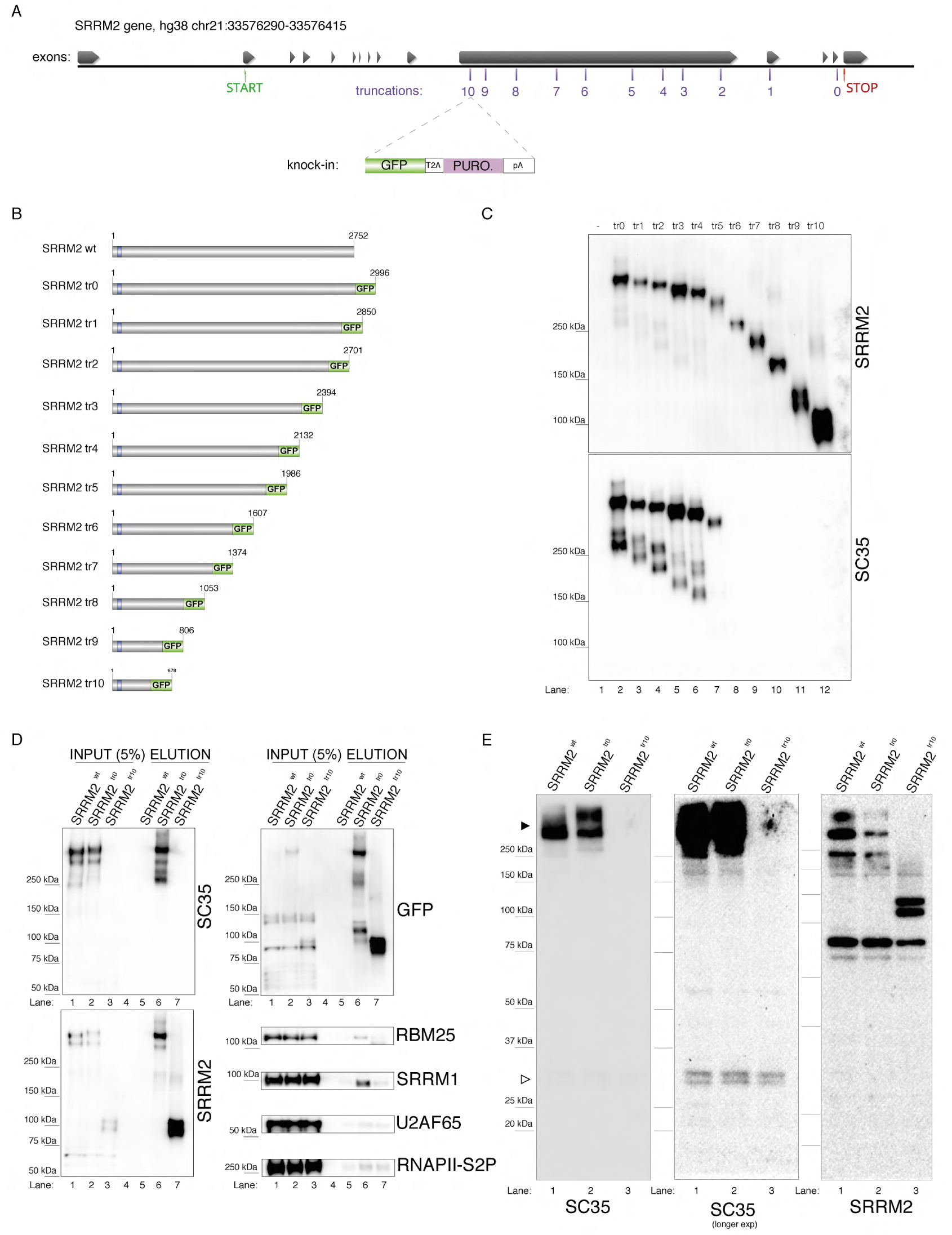
Endogenous truncating mutations of SRRM2 prove mAb SC35 as an SRRM2 antibody. **(A)** The strategy for the CRISPaint generated endogenous truncating mutations (0-to-10) accompanied by the TagGFP2 (depicted as GFP for simplicity) fusion are shown. **(B)**The size of SRRM2 truncated GFP fusion proteins are displayed. **(C)** Protein purified using a GFP-trap pull-down from lysates of corresponding stable HAP1 cell lines carrying the truncated SRRM2 alleles are run on PAGE. Western blotting of SRRM2 using an antibody generated against the common N-terminus is used to show the amount of loaded protein on the gel. SC35 blot shows a significant reduction in signal intensity of SRRM2-tr5 and a complete loss of signal from SRRM2-tr6 to tr10. **(D)** GFP-trap pull-down performed on lysates from wild-type, tr0 and tr10 HAP1 cells enrich for SRRM2 in tr0 cells, indicating the GFP tagged allele is specific to SRRM2 and is detected by also SC35 blot (Lanes 1 and 2 inputs compared to Lanes 5 and 6 on the upper and lower left-side blots). SRRM2 also co-purifies two other NS associated proteins; SRRM1 and RBM25 (Lane 6 on lower right-side blots). SRRM2-tr10 is not detected by SC35 but the pull-down efficiency (Lane 7 on upper left-side blot) and loading is validated by SRRM2 (Lanes 3 and 7 on lower left-side blot) and GFP blots (Lanes 3 and 7 on upper left-side blot). **(E)** Total cell lysates from wild-type, tr0 and tr10 HAP1 cells are run on 4-12% PAGE gel and blotted with SC35 reveal the high molecular weight (∼300kDa) as the most intense band and the absence of signal in tr10 cell lines validates that this band represents SRRM2 (filled arrow head). Longer exposure of the blot reveals a weak cross reactivity with a 35 kDa protein, most likely to be SRSF7, around 35kDa (empty arrow head).

In order to assess the efficiency and the specificity of the GFP-pull-down, we used a wild-type lysate without any GFP insertion, together with lysates made from SRRM2^tr0^ and SRRM2^tr10^ cells, which served as the negative control, positive control and the deepest truncation (tr10) we generated, respectively. The immunoblot with mAb SC35 once again clearly shows that near-full-length SRRM2^tr0^ is recognized by mAb SC35 to the same extent as the SRRM2 polyclonal antibody, while SRRM2^tr10^ is not detected by mAb SC35 at all but strongly with SRRM2 polyclonal antibody (Fig. 2D). These blots also show that full-length SRRM2 co-purifies SRRM1 and to a lesser extent RBM25, while both interactions are severely compromised in SRRM2^tr10^. Furthermore, the absence of any signal in SRRM2^tr10^ input lane probed with mAb SC35 (Fig. 2D, top left lane 3), and the emergence of a shorter ∼100kDa protein in the complete absence of a ∼300kDa signal in the SRRM2 blot (Fig. 2D, bottom left, lane 3) shows that SRRM2^tr10^ cells have a homozygous insertion of the GFP construct, which was also confirmed by genotyping PCR (Supplementary Fig. 3D). This result further indicates that the large IDRs of SRRM2 are not essential for cell viability, at least in HAP1 cells.

These results can be puzzling, since we first show that mAb SC35 specifically recognizes a 35 kDa band which we reveal to be SRSF7 (Fig. 1), but later, in a separate set of experiments, we also show that mAb SC35 specifically recognizes a ∼300 kDa band, which we reveal to be SRRM2 (Fig. 2), while the original study describing mAb SC35 reports a single 35 kDa band recognized by mAb SC35 on immunoblots (6). The solution to this conundrum presented itself in the form of altering the immunoblotting technique. Using whole-cell extracts prepared from wild-type cells, together with SRRM2^tr0^ and SRRM2^tr10^ cells, in a gel system where we can interrogate both small and large proteins simultaneously, we were able to detect both SRRM2 and SRSF7 on the same blot (Fig. 2E). These blots prove that the ∼300 kDa band is indeed SRRM2, since it completely disappears in SRRM2^tr10^ lysates (which is accompanied by the appearance of a ∼100 kDa band in SRRM2 blots) while the much fainter 35 kDa band corresponding to SRSF7 (Fig. 1) remains unaltered.

These experiments provide strong support for our hypothesis that the main target of SC35 mAb is SRRM2, a protein proven to be part of spliceosomes, against which this antibody was raised, and suggests that a cross-reactivity towards SRSF7, likely in combination with immunoblotting techniques not suitable to detect large proteins (36), obscured this fact for more than two decades.

### SRRM2 is the primary target of mAb SC35 in immunofluorescence stainings

mAb SC35 is typically used as an antibody in immunofluorescence experiments that reveals the location of nuclear speckles in mammalian cells (3). In light of the evidence presented here, it can be assumed that mAb SC35 primarily stains SRRM2 in immunofluorescence stainings, as in immunoblotting experiments. In order to test if this is indeed the case, we took advantage of the SRRM2^tr10^ cells. These cells are viable and express a severely truncated SRRM2 that is not recognized by mAb SC35 on immunoblots (Fig. 2).

SRRM2^tr10^ cells, together with SRRM2^tr0^ cells serving as a control, were stained with antibodies against various nuclear speckle markers, including mAb SC35 (Fig. 3A). These results show that SC35 signal virtually disappears in SRRM2^tr10^ cells, while other markers of nuclear speckles, such as SON, SRRM1 and RBM25 appear unaltered, ruling out a general defect in nuclear speckles (Fig. 3A). As an additional control, we also mixed SRRM2^tr10^ cells with SRRM2^tr0^ cells together before formaldehyde fixation, and repeated the antibody stainings, in order to be able to image these two cell populations side-by-side. SRRM2^tr10^ and SRRM2^tr0^ cells are easily distinguished from each other since the latter show a typical nuclear speckle staining whereas the former has a more diffuse, lower intensity GFP signal. These images clearly show that mAb SC35, obtained from two separate vendors, no longer stains nuclear speckles or any other structure in SRRM2^tr10^ cells (Fig. 3B,C).

**Fig. 3.**
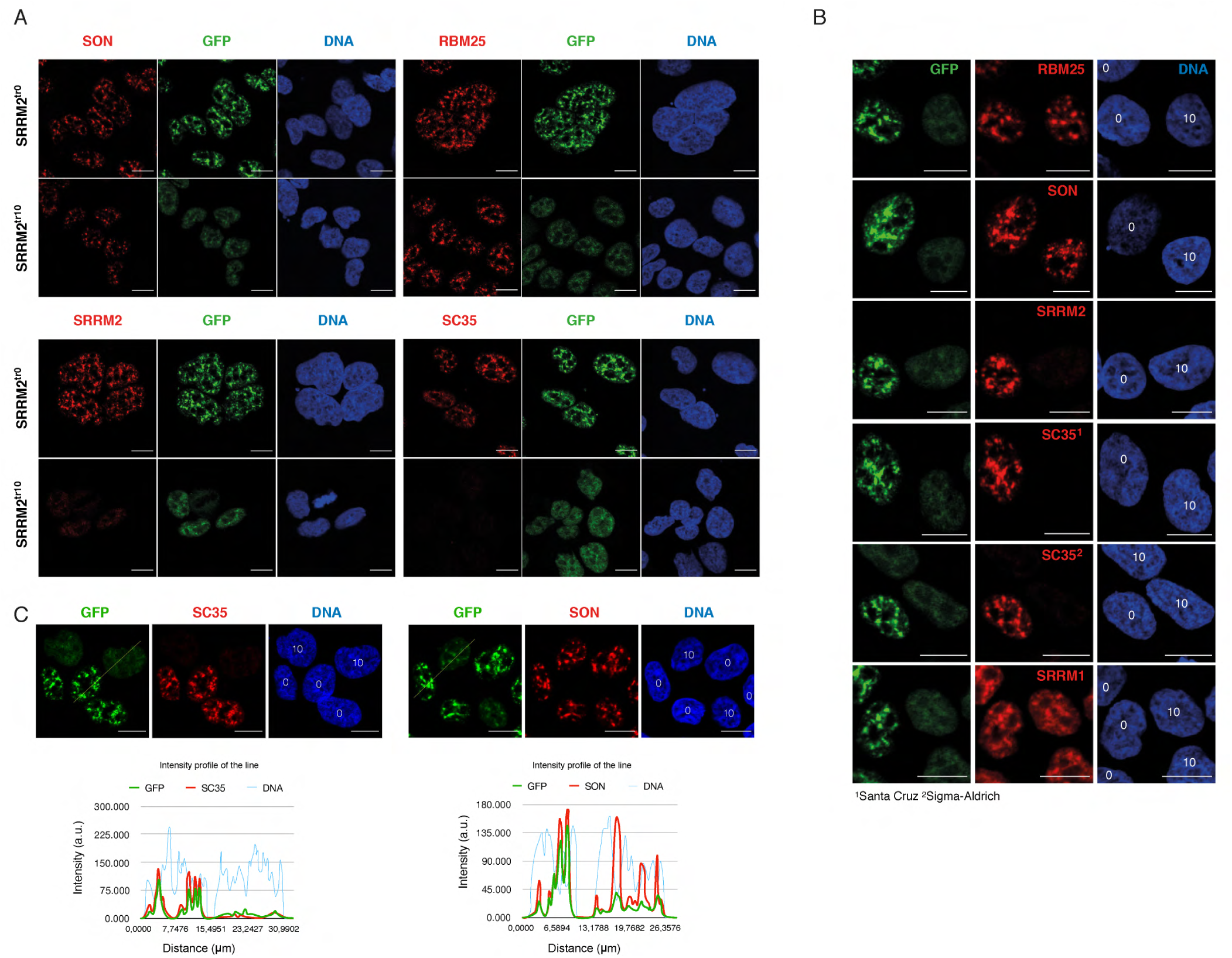
SRRM2 truncation#10 leads to loss of SC35 domains but not NS. **(A)** SON and RBM25 antibodies are used as NS markers for IF analysis of both SRRM2^tr0^ and SRRM2^tr10^ HAP1 cells. No significant impact on the formation of NS in SRRM2^tr10^ cells in comparison to SRRM2^tr0^ cells is observed. Lack of signal for SC35 in SRRM2^tr10^ cells validates SC35 as an SRRM2 antibody. **(B)** The SRRM2^tr0^ and SRRM2^tr10^ cells are plated together before the IF protocol is performed and the GFP signal intensity as well as SC35 staining are used to distinguish SRRM2^tr10^ cells from SRRM2^tr0^ HAP1 cells. The DNA stain marking the nuclei are annotated with “0” or “10” on top to indicate the corresponding cell line. **(C)** The SRRM2^tr0^ and SRRM2^tr10^ HAP1 cells are imaged side by side and a line is drawn to quantify the signal intensity across two cell lines. The intensity profile of the lines show dramatically reduced signal for SC35 between SRRM2^tr0^ and SRRM2^tr10^ cells, whereas similar signal intensities for SON and DNA in SRRM2^tr0^ cells is observed. Scale bars = 10µm.

Taken together, our results show that mAb SC35, which was raised against a spliceosomal extract, was most likely raised against SRRM2, a ∼300kDa protein that, unlike SRSF2 or SRSF7 is present in spliceosomes of both in yeast and humans. We show that mAb SC35 directly recognizes SRRM2 between amino acids 1,360 - 1,884, and that the main signal from mAb SC35 corresponds to SRRM2 both in immunoblots and immunofluorescence images. It is interesting to note that this is not the first time an antibody is serendipitously raised against SRRM2 and was later discovered to recognize SRRM2 only after the fact: In 1994, Blencowe et al. reported three murine monoclonal antibodies, B1C8, H1B2 and B4A11 which were raised against nuclear matrix preparations (8). All three antibodies showed extensive co-localization with nuclear speckles, although a co-localization between mAb SC35 and B4A11 could not directly be assessed since both mAb SC35 and B4A11 are reported to be IgG mAbs. In a separate work, Blencowe et al. showed that B4A11 is an antibody against SRRM2 (25), suggesting that SRRM2 is present both in spliceosomal purifications and nuclear matrix preparations.

### NS formation requires SON and full-length SRRM2

Our results have broad implications with respect to the biology in and around nuclear speckles. One of the primary culprits in the so-called “reproducibility crisis” in natural sciences is considered to be mischaracterized antibodies (37), which led to initiatives to validate them appropriately (38), and large consortia such as ENCODE (39) publishes specific guidelines for the characterization antibodies that are used to generate data pertinent to the ENCODE project (40).

It is therefore reasonable to suspect that the dissonance between observations made with mAb SC35 and subsequent observations made with SRSF2 reagents could have led to misinterpretation of vast amounts of primary data. It is beyond the scope of this work to review each and every study that has used this antibody in the last 30 years, however, since both *SRSF2* and *SRRM2* are clinically important genes (41–47) and SRRM2 gene is highly to intolerant to loss-of-function mutations in human populations (expected/observed ratio for SRRM2 is 6%, median expected/observed ratio for all gene variants is 48%) (48), the opportunity cost incurred when results generated with mAb SC35 have been misinterpreted must be carefully considered.

For example, GSK-3, a protein kinase implicated in Alzheimer’s disease (49), has been shown to phosphorylate SRSF2 (50). This study reported that inhibition of GSK-3 leads to spherical NS in cortical neurons, which was interpreted as SRSF2 accumulating at NS upon GSK-3 inhibition. Based on this assumption, a cerebral cortex lysate was immunoprecipitated with mAb SC35, and a 35 kDa band was shown to be phosphorylated by recombinant GSK-3, whereas neither SRSF7 nor SRRM2 were pursued as potential candidates. Interestingly, a more recent study reported that SRRM2 is a target of GSK-3 (51). In another study, mAb SC35 has been used as a reporter for phosphorylated SRSF2 in immunohistochemical analysis of tissue samples obtained from patients with Non Small Cell Lung Carcinoma (52), and was reported to be upregulated in cancer tissues compared to normal tissues. SRRM2’s potential role in these syndromes therefore remains unexplored.

In a more non-clinical setting, recent microscopy work suggests that RNA polymerase II, through its intrinsically disordered CTD, switches from transcriptional condensates to NS during progression from initiation to productive elongation (53), which is in line with recent work that links NS to augmented gene expression (5). In this particular work, initial observations made with the SC35 mAb have been followed by fluorescently tagged SRSF2 protein for live-cell imaging analysis and condensates formed by SRSF2 are used as a surrogate for splicing condensates *in vitro*. Furthermore, ChIPseq profiles generated with mAb SC35 were interpreted as SRSF2 occupancy *in vivo*, which was shown to be enriched at the 3’-ends of genes, together with serine-2 phosphorylated RNAPII. SRRM2’s potential role in these processes remain unknown. In another study, loss of mAb SC35 signal in cells treated with siRNAs against SRRM2 has been interpreted to be a sign of NS disassembly (32), and based on this knowledge, a more recent study interrogated the 3D conformation of the mouse genome in SRRM2-depleted cells using Hi-C (54), results of which are interpreted to be a consequence of NS disruption. A recent super resolution microscopy study made use of the mAb SC35 to show that SRSF2 is at the core of NS together with SON, and remains at the core after depletion of SON (55). In this study, a minimalist computational model was used to model the distribution of five NS associated factors (SON, SRSF2 on account of SC35 mAb stainings, MALAT1, U1 and U2B”) which could have benefited from the knowledge that SC35 mAb stains SRRM2 rather than SRSF2, considering that pairwise interactions between SRRM2 and other components would be drastically different to SRSF2’s potential interactions due to the extensive IDRs of SRRM2 compared to SRSF2. This study also highlights the fact that a crucial aspect of nuclear speckle biology, namely whether NS are nucleated by specific factors or not, remains an open question. In contrast, many membrane-less organelles have been shown to depend on a small number of factors which act as scaffolds or nucleation points for their formation. Paraspeckles for instance require lncRNA NEAT1, without which paraspeckles do not form (56), Cajal bodies are disrupted or disappear in the absence of COILIN, SMN, FAM118B or WRAP53 (57, 58), and PML bodies are nucleated by PML (59). In the same vein, SRRM2 has been suggested to be essential for the formation of nuclear speckles (32), however, this idea was based on the disappearance of mAb SC35 signal in cells transfected with siRNAs against SRRM2, which is the expected result taking the evidence presented here into account, but does not prove that SRRM2 is essential or important for NS formation. Other candidates that were put forward as essential or important for the formation of NS include lncRNA MALAT1 (60, 61), SRSF1 (17), PNN and SON (20, 55, 62), all of which, with the exception of PNN, lead to the formation of “collapsed” speckles rather than a bulk release of NS-associated factors and their diffusion into the nucleoplasm, which would indicate a true loss of NS. Depletion of PNN was shown to either lead to “collapsed” speckles (63) or to loss of NS altogether, but under conditions that also lead to degradation of all SR-proteins tested in that particular study (64). To our knowledge, NS could only be successfully dissolved by overexpression of CLK1/STY kinase, which phosphorylates SR-proteins (15), DYRK3, another protein kinase that can dissolve multiple membraneless bodies (14), overexpression of PPIG, a peptidyl-proline isomerase (65) or more recently by overexpression of TNPO3, which is an import factor that binds to phosphorylated SR-residues (66). Such observations and lack of evidence to the contrary, led to the idea that NS formation happens through stochastic self-assembly of NS-associated factors, without the need for an organizing core (3, 16, 17).

During this work, we noticed the remarkable size difference between human SRRM2 protein (2752 a.a.), and its unicellular counterparts *S. cerevisiae* Cwc21 (133 a.a), *S. pombe* Cwf21 (293 a.a) and *T. Brucei* U5-Cwc21 (143 a.a). Moreover, while all three proteins share a conserved N-terminus, which interact with the spliceosome, the serine and arginine rich extended C-terminus of human SRRM2 is predicted to be completely disordered (Supplementary Fig. 3B). Intrigued by this observation, we compiled all metazoan protein sequences of SRRM2, together with SRRM1, RBM25, PNN, SON, PRPF8 and COILIN, and analyzed their size distributions (Fig. 4A). This analysis confirmed that, unlike SRRM1, RBM25, PNN, PRPF8 or coilin, SRRM2 indeed has a very broad size distribution within metazoa (Supplementary Fig. 4, Supplementary Fig. 5). Strikingly, SON follows this trend with orthologues as small as 610 a.a in the basal metazoan sponge *A. queenslandica*, and as large as 5,561 a.a in the frog *X. tropicalis*. Increase in protein size appears to involve IDR extensions, especially for SRRM2, but also for SON (Fig. 4B, Supplementary Fig. 4 and Supplementary Fig. 5), suggesting a role in LLPS-mediated condensate formation, which was shown to be the case for both SRRM2 (14) and SON (67) in living cells.

**Fig. 4.**
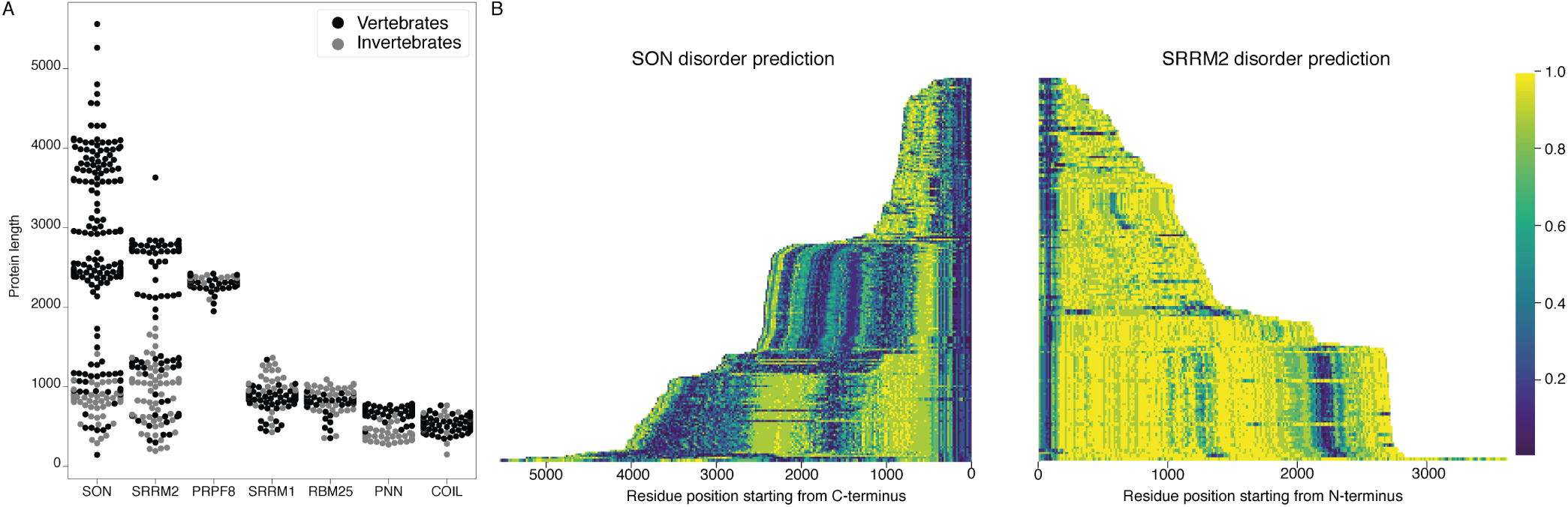
SON and SRRM2 are rapidly evolving and largely disordered proteins. **(A)** The size distribution of SON and SRRM2 is highly variable across metazoan species with a mean length of 2227.9 a.a. and SD of 1149.5 for SON and a mean length of 1928.6 a.a. and SD of 919.3 for SRRM2. The lengths of other NS associated proteins are less variable with a mean length of 895.1 a.a and SD of 104.2 for SRRM1; mean length of 835.4 a.a. and SD of 77.9 for RBM25; mean length of 652.8 a.a. and SD of 118.7 for PNN, mean length of 2332.8 a.a. SD of 40.8 for PRP8. **(B)** The disorder probability of SON and SRRM2 is predicted using the MobiDB-Lite algorithm, which shows an increase of disordered content with the increase of protein length for SRRM2, and to some extent, SON. The SON and SRRM2 graphs plotted side-by-side do not correspond to the same species, for a phylogeny resolved version of this graph see Supplementary Fig. 4 and for the alternative algorithm (IUPred2A) see Supplementary Fig. 5. The color is scaled from dark blue to yellow indicating a decrease in order as the value approaches 1.0 (yellow).

Putting together the observation that places SON and SRRM2 at the center of NS ((55), with the interpretation that SC35 stains SRRM2 in their microscopy work), the presence of SRRM2 at the center of collapsed speckles in SON knock-down experiments, and the peculiar variation in the sizes of SON and SRRM2 during evolution involving gain of IDRs, we hypothesize that SON, together with SRRM2 are essential for NS formation, such that SRRM2 continues to serve as a platform for NS-associated proteins in SON-depleted cells.

In order to test this hypothesis, we used the SRRM2^tr0^ and SRRM2^tr10^ cells as a model, which allowed us to simultaneously detect SON, SRRM2 and an additional NS marker in the same cell. We chose SRRM1, which is used as a marker for NS in immunofluorescence experiments (8, 9, 14, 25, 68) and located at ICGs in electron microscopy experiments (69); RBM25, which is one of two recommended factors to mark NS by the Human Protein Atlas (70) (the other being SRRM2),localizes to NS through its RE/RD-rich mixed-charge domain (71) that was recently shown to target proteins to NS (72) and PNN, which localizes to nuclear speckles in human cells (63–65, 73).

As reported previously (20, 55, 62), depletion of SON leads to collapsed speckles in SRRM2^tr0^ cells, with SRRM2, SRRM1, PNN and RBM25 localizing to these spherical NS to different extents (Fig 5A, Fig. 5B, Supplementary Fig. 8, compare SRRM2^tr0^ cells, control vs SON siRNA treatment). In SRRM2^tr10^ cells on the other hand, where the truncated SRRM2 has a significant nucleoplasmic pool already in control siRNA treated cells, depletion of SON leads to a near-complete diffusion of truncated SRRM2, which is followed by RBM25, SRRM1 and PNN. Using ilastik and CellProfiler, we quantified the signal detected in NS, and compared it to signal detected in the entire nucleus for each cell in every condition for each protein investigated (Supplementary Fig. 6 and Supplementary Fig. 7). These results show that truncated SRRM2 shows reduced NS localization (Fig. 5C, right), while RBM25, SRRM1 (Fig. 5B) and PNN (Supplementary Fig. 8A) are localized at NS to a similar extent in SRRM2^tr0^ and SRRM2^tr10^ cells, although with a broader distribution in SRRM2^tr10^ cells. Depletion of SON in SRRM2^tr0^ leads to a significant reduction in NS localization for all proteins, verifying SON’s importance for NS formation. Depletion of SON in SRRM2^tr10^ cells, however, leads to a more dramatic loss of NS localization for all proteins (Fig. 5 and Supplementary Fig. 8), underscoring the essential role of SRRM2’s extended IDR in the formation of NS, especially in SON-depleted cells. Number of Cajal bodies, determined by COILIN staining, remains unaltered in all conditions (Supplementary Fig. 8B).

**Fig. 5.**
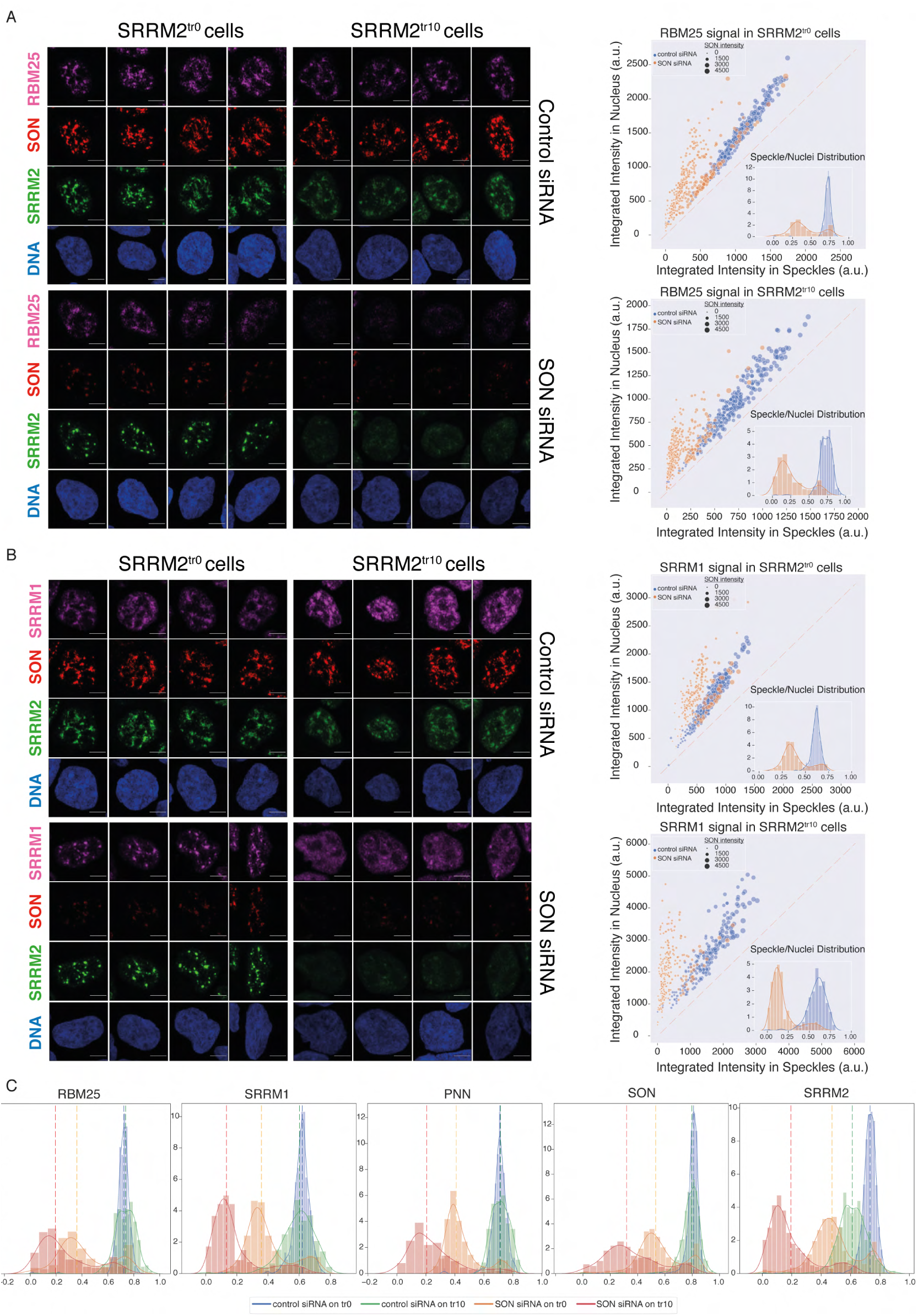
SON and SRRM2 form NS in human cells. **(A)** RBM25 IF signal is shown for four individual cells in each siRNA treatment (control or SON siRNA) in SRRM2^tr0^ and SRRM2^tr10^ HAP1 cells. The NS localization of RBM25 is severely reduced upon SON knock-down in SRRM2^tr0^ cells, and completely lost upon SON knock-down in SRRM2^tr10^ cells. The quantification of the RBM25 signal within the nucleus is plotted against the RBM25 signal within NS (right panel) using ilastik to train detection of NS and CellProfiler for quantification on 10 imaged fields with a 63X objective (in SRRM2^tr0^ cells control n= 329, SON-KD n=422; in SRRM2^tr10^ cells control n= 329, SON-KD n=402). Each circle represents a cell and the size of the circles is proportionate to the signal intensity of SON. Inset shows the distribution of the ratio of signal detected in NS over signal detected in the nucleus of each cell. **(B)** SRRM1 IF signal is shown for four individual cells in each siRNA treatment (control or SON siRNA) in SRRM2^tr0^ and SRRM2^tr10^ HAP1 cells. The NS localization of SRRM1 is reduced in SON knock-down in SRRM2^tr10^ cells and lost upon SON knock-down in SRRM2^tr10^ cells. The quantification of the SRRM1 signal within the nucleus is plotted against the SRRM1 signal within NS (right panel) using ilastik to train detection of NS and CellProfiler for quantification on 10 imaged fields with 63X objective (in SRRM2^tr0^ cells control n=494, SON-KD n=229; in SRRM2^tr10^ cells control n=225, SON-KD n=247). Inset shows the distribution of the ratio of signal detected in NS over signal detected in the nucleus of each cell. **(C)** Distribution plots showing the ratio of signal detected in NS over signal detected in the nucleus of each cell, in each condition. The dashed line indicates the median ratio in each condition. See Supplementary Fig. 8A for a full version of this analysis for PNN. Scale bars = 5µm.

Next, to independently verify these observations, we knocked-down SON and SRRM2, individually and simultaneously in HEK293 cells where we endogenously tagged SRRM2 with TagGFP2 at the C-terminus with the same reagents used to create SRRM2^tr0^ HAP1 cells. Similar to the HAP1 model, depletion of SON alone leads to collapsed NS where SRRM2, RBM25, PNN and SRRM1 localize to spherical NS to some extent but with a significant non-NS pool in the nucleus (Supplementary Fig. 9). Depletion of SRRM2 alone also leads to delocalization of PNN, SRRM1 and RBM25 from NS, but not to the extent seen with SON depletion. Co-depletion of SON and SRRM2 lead to nearcomplete delocalization of all proteins investigated, mirroring the results obtained from the HAP1 model (Supplementary Fig. 9A,B,C and D). These results cannot be explained by reduced protein stabilities, as none of the proteins except for SON and SRRM2 show significant changes in their amounts as judged by immunoblotting (Supplementary Fig. 9E). Finally, co-depletion of SON together with SRRM1 or RBM25 does not lead to diffusion of spherical NS marked by SRRM2, indicating that SRRM2 has a unique role in NS formation and cannot be substituted by other NS-associated factors (Supplementary Fig. 10).

## Conclusions

Taken together, our results show that a widely-used monoclonal antibody to mark nuclear speckles, SC35 mAb, was most likely raised against SRRM2 and not against SRSF2 as it was initially reported. We speculate that this mischaracterization hindered the identification of the core of nuclear speckles, which we show to consist of SON and SRRM2. We found that these two factors, unlike other splicing related proteins analyzed, have gone through a remarkable length extension through evolution of metazoa over the last ∼0.6-1.2 billion years, mostly within their IDRs which are typically involved in LLPS and formation of biomolecular condensates. The exact mechanism of NS formation by SON together with SRRM2, and the evolutionary forces that led to the dramatic changes in their lengths remain to be discovered.

## Supporting information

Summary Figure

## ACKNOWLEDGEMENTS

We thank Florian Heyd, Denes Hnisz and Alexander Meissner for critical reading of the manuscript and helpful suggestions. We thank the Mass Spectrometry Facility at MPI-MG, especially Beata Lukaszewsa-McGreal, members of the Microscopy and Cryo-Electron Microscopy Facility at MPI-MG, especially Thorsten Mielke, Beatrix Fauler and René Buschow. We thank Christina Riemenschneider (Laboratory of Alexander Meissner) and Mirjam Arnold (Laboratory of Andreas Mayer) for sharing reagents, Prof. Feng Zhang and Prof. Veit Hornung for sharing their plasmids via Addgene, developers and maintainers of Open-source software, especially ilastik, CellProfiler, STRINGdb, Python, SciPy, NumPy, Matplotlib, Jupyter Lab, seaborn and Dr. Ricardo Henriques for making this bioRxiv template publicly available. Initial cell culture work was only possible thanks to the help of Sebastiaan Meijsing Group. Research in the laboratory of T.A. is funded by the Max-Planck Research Group Leader program.

## Author Contributions

This study was perceived and designed by IAI and TA. IAI carried out most of the experiments with assistance from CS and KL. MM carried out the phylogenetic analysis under the supervision of IAI and TA. DM carried out the Mass Spectrometry and analysis. IAI and TA wrote the manuscript.

## Materials and Methods

### Cell culture and generation of stable cell lines

Flp-In T-REx HEK293 (Thermo Fisher Scientific Catalog Number: R78007) cells were kept according to manufacturer’s recommendations. The cells were cultured in DMEM with Glutamax supplemented with Na-Pyruvate and High Glucose (Thermo Fisher Scientific Catalog Number: 31966-021) in the presence of 10% FBS (Thermo Fisher Scientific Catalog Number: 10270106) and Penicillin / Streptomycin (Thermo Fisher Scientific Catalog Number: 15140-122). Before the introduction of the transgenes cells were cultured with a final concentration of 100µg/ml zeocin (Thermo Fisher Scientific Catalog Number: R250-01) and 15µg/ml blasticidin(Thermo Fisher Scientific Catalog Number: A1113903). To generate the stable cell lines pOG44 (Thermo Fisher Scientific Catalog Number: V600520) was co-transfected with pcDNA5/FRT/TO (Thermo Fisher Scientific Catalog Number: V652020) containing the gene of interest (GOI are SRSF1 to 12 in this case) in a 9:1 ratio. Cells were transfected with Lipofectamine 2000 (Thermo Fisher Scientific Catalog Number: 11668019) on a 6-well plate format with total 1µg DNA (i.e. 900ng of pOG44 and 100ng of pcDNA5/FRT/TO+GOI) according to the transfection protocol provided by the manufacturer. 24 hours after the transfection cells were split on 3 wells of a 6-well plate at 1:6, 2:6 and 3:6 dilution ratios to allow efficient selection of Hygromycin B (Thermo Fisher Scientific Catalog Number: 10687010). The Hygromycin selection was started at the 48 hours after transfection time point with a final concentration of 150µg/ml and refreshed every 3-4 days until the control non-transfected cells on a separate plate were completely dead (takes approximately 3 weeks from the start of transfection until the cells are expanded and frozen). Induction of the transgene was done over-night with a final concentration of 0.1µg/ml doxycycline. The cells were validated by performing immunofluorescence by FLAG antibody and western blotting of nuclear and cytoplasmic fractions.

Human HAP1 parental control cell line was purchased from Horizon (Catalog Number: C631) and cultured according to the instructions. The cells were cultured in IMDM (Thermo Fisher Scientific Catalog Number: 12440-053) in the presence of 10% FBS (Thermo Fisher Scientific Catalog Number: 10270106) and Penicillin / Streptomycin (Thermo Fisher Scientific Catalog Number: 15140-122). CRISPaint constructs (see Supplementary Table 1 for the list of sgRNAs). Cells were co-transfected with 3 plasmids according to the CRISPaint protocol. Cas9 and sgRNA are provided by same plasmid in 0.5µg final amount, Frame selector plasmid (depending on the cut site selector 0, +1 or +2 had to be chosen) is also in 0.5µg final amount, the TagGFP2_CRISPaint plasmid was provided at a 1µg final amount. Therefore the total 2µg DNA was transfected into cells on 6-well plate format using Lipofectamine 2000. 24 hours after the transfection the cells were expanded on 10cm culture plates to allow efficient Puromycin (Thermo Fisher Scientific Catalog Number: A1113803) selection. The Puromycin selection is provided in the tag construct and is driven by the expression from the gene locus (in this case the human *SRRM2* gene locus). Puromycin selection was started at 48 hours after transfection at 1µg/ml final concentration and was refreshed every 2 days and in total was kept for 6 days. After the colonies grew to a visible size the colonies were picked by the aid of fluorescence microscope EVOS M5000. PCR screening of the colonies was performed using genotyping oligos listed in Supplementary Table 1 using Quick Extract DNA Extraction Solution (Lucigen Catalog Number: QE09050) according to manufacturer’s protocol in a PCR machine and DreamTaq Green Polymerase (Thermo Fisher Scientific Catalog Number: K1081) using 58°C annealing temperature and 1 minute extension time.

SRSF7-FKBP12 ^F36V^ knock-in cells were generated in HEK293T cells (ordered from ATCC, CRL3216 and cultured according to the protocol provided) by co-transfecting the sgRNA, Frame selector and mini-circle constructs prepared according to the CRISPaint protocol using Lipofectamine 2000 on a 6-well plate format. This time we used two separate tag donor plasmids to increase the chances of obtaining homozygous clones. The constructs were identical except for the selection antibiotic. Cells are expanded on 10cm culture plates 24 hours after transfection. At 48 hours after transfection the double selection was initiated. One allele was selected by Puromycin at 1µg/ml final concentration, whereas the other allele was selected by blasticidin at 15µg/ml final concentration for 6 days in total. After the removal of selection cells were kept on the same plate until there were big enough colonies. Colonies were picked under a sterile workbench and screened for homozygosity using western blotting with SRSF7 antibody. The degradation of tagged-SRSF7 was induced by adding dTAG7 reagent at a final 1µM concentration and keeping for 6 hours.

SRRM2^tr0^-GFP Flp-In TREx HEK293 cells were generated using the same strategy as described above for HAP1 cells. Upon Puromycin selection cells were used as a pool (without sorting or colony picking) in immunofluorescence experiments.

Cell lines are regularly checked for the absence of Mycoplasma using a PCR based detection kit (Jena Biosciences PP-401).

### siRNA transfections

Prior to the seeding of cells, the round glass 12mm coverslips are coated with poly-L-Lysine hydrobromide (Sigma P9155) for HEK293 cells. The coating is not necessary for the imaging of HAP1 cells. For 1 day of knock-down 40,000 cells are plated on coverslips placed into the wells of 24-well plates on the day before the siRNA transfections. Pre-designed silencer select siRNA (Ambion) are ordered for SRRM2 (ID: s24004), SON (ID: s13278), SRRM1 (ID: s20020) and RBM25 (ID: s33912). Negative control #1 of the silencer select was used for control experiments. 5nM (for double transfections) or 10nM (for single transfections) of each siRNA is forward transfected using Lipofectamine RNAiMAX Reagent (Thermo Fisher Scientific Catalog Number: 13778075) according to manufacturer’s instructions. The cells are fixed for imaging 24 hours after transfection.

### Immunofluorescence and imaging

#### Sample preparation

Cells on coverslips were washed with PBS and crosslinked with 4% paraformaldehyde in PBS (Santa Cruz Biotechnology, sc-281692) for 10 minutes at room temperature, and washed three times with PBS afterwards. Permeabilization was carried out with 0.5% Triton-X in PBS, 10 minutes at RT. Cells were washed twice with 0.1% Triton-X in PBS and blocked with 3% BSA (constituted from powder BSA, Roche Fraction V, sold by Sigma Catalog Number: 10735078001) in PBS for 30 minutes at RT. Primary antibodies were diluted in 3% BSA in PBS, and cells were incubated with diluted primary antibodies for ∼16hrs at 4°C in a humidified chamber. Cells were then washed three times with 0.1% Triton-X in PBS and incubated with fluorescently labelled secondary antibodies, diluted 1:500 in 3% BSA for 1hr at RT, and washed three times with 0.1% Triton-X in PBS. To counterstain DNA, cells were incubated with Hoechst 33258 (1µg/mL, final) for 5 minutes at RT, and washed once with PBS. Coverslips are briefly rinsed with distilled water and mounted on glass slides using Fluoromount-G® (SouthernBiotech, 0100-01) and after a few hours, sealed with CoverGrip™ (Biotium, #23005) and left in a dark chamber overnight before imaging.

#### Antibodies

COIL (Cell Signaling Technology, D2L3J, 14168), FLAG-M2 (Sigma, F3165), PNN (Abcam, ab244250), RBM25 (Sigma, HPA070713-100UL), SC-35 (Santa Cruz Biotechnology, sc-53518), SC-35 (Sigma, S4045), SON (polyclonal rabbit, Sigma, HPA023535), SON (monoclonal mouse, Santa Cruz sc-398508), SRRM1 (Abcam, ab221061), SRRM2 (Thermo Fisher Scientific, PA5-66827).

#### Imaging

Images were acquired with a Zeiss LSM880 microscope equipped with an AiryScan detector, using the AiryScan Fast mode with the Plan-Apochromat 63x/1.40 Oil DIC M27 objective. The dimensions of each image was 134µm x 134µm x 4µm (Width x Height x Depth), 20 z-stacks were acquired for each image with a step size of 200nm. Maximum Intensity Projections were created using Zen software (Zeiss) and used for further analysis.

#### Analysis

Nuclear speckle identification, segmentation and intensity calculations were carried out using ilastik and CellProfiler. Briefly, eight images were used to train a model that demarcates NS using ilastik (v.1.3.3post2). CellProfiler was then used to segment nuclei and NS using the probability maps created for each image by ilastik. The data was then analyzed in a Jupyter Lab environment using pandas, SciPy, NumPy and plotted with matplotlib and seaborn. Raw imaging data, models used to train the images, CellProfiler pipelines and Jupyter notebooks will be available.

### Mass Spectrometry. Sample preparation

Pierce MS-Compatible Magnetic IP Kit (Protein A/G) (Thermo Fisher Scientific, Catalog Number: 90409) was used to prepare samples for mass-spectrometry according manufacturer’s instructions, where approximately 15 million HAP1 cells (∼80% confluent 15cm dishes) were used per IP. Briefly, HAP1 cells were trypsinized, washed with ice-cold PBS and re-suspended with 500µL of ‘IP-MS Cell Lysis Buffer’ which was supplemented with 1x cOmplete™ Protease Inhibitor Cocktail (Roche, 11697498001) and 1x PhosSTOP™ (Roche, 4906845001). Cells were then homogenized using a Bioruptor® Plus sonifier (30s ON, 30s OFF, 5 cycles on HI). Remaining cellular debris was removed by centrifugation at 21.130 rcf for 10 minutes at 4°C, supernatants were transferred to fresh tubes. 2.5µL of SC35 mAb (Sigma-Aldrich, S4045) and 25µL of control IgG1 (Santa Cruz, sc-3877) was used for the SC35 and control IP samples (3 each), respectively and immune-complexes are allowed to form overnight (∼16hrs) in the cold-room (∼6°C) with end-to-end rotation. Next morning, lysates were incubated with 25µL of Protein A/G beads for 1hr in the cold-room, the beads were then washed with 500µL of ice-cold 50mM Tris.Cl pH 7.4, 100mM NaCl, 0.1% Tween-20. The beads were resuspended with the same buffer supplemented with RNaseI (Ambion, AM2295, final concentration 0.02 U/µL) and incubated at 37°C for 5 minutes. The beads were then washed 3 times with ‘Wash A (+10mM MgCl2)’ and twice with ‘Wash B’ buffer.

### On beads digest and mass spectrometry analysis

The buffer for the three SC35 samples and controls was exchanged with 100 µl of 50 mM NH_4_HCO_3_. This was followed by a tryptic digest including reduction and alkylation of the cysteines. Therefore, the reduction was performed by adding tris(2-carboxyethyl)phosphine with a final concentration of 5.5 mM at 37°C on a rocking platform (500 rpm) for 30 minutes. For alkylation, chloroacetamide was added with a final concentration of 24 mM at room temperature on a rocking platform (500 rpm) for 30 minutes. Then, proteins were digested with 200 ng trypsin (Roche, Basel, Switzerland) shaking at 600 rpm at 37°C for 17 hours. Samples were acidified by adding 2.2 µL 100% formic acid, centrifuged shortly, and placed on the magnetic rack. The supernatants, containing the digested peptides, were transferred to a new low protein binding tube. Peptide desalting was performed according to the manufacturer’s instructions (Pierce C18 Tips, Thermo Scientific, Waltham, MA). Eluates were lyophilized and reconstituted in 11 µL of 5% acetonitrile and 2% formic acid in water, briefly vortexed, and sonicated in a water bath for 30 seconds prior injection to nano-LC-MS/MS.

### LC-MS/MS Instrument Settings for Shotgun Proteome Profiling and Data Analysis

LC-MS/MS was carried out by nanoflow reverse-phase liquid chromatography (Dionex Ultimate 3000, Thermo Scientific) coupled online to a Q-Exactive HF Orbitrap mass spectrometer (Thermo Scientific), as reported previously (74). Briefly, the LC separation was performed using a PicoFrit analytical column (75 µm ID × 50 cm long, 15 µm Tip ID; New Objectives, Woburn, MA) in-house packed with 3-µm C18 resin (Reprosil-AQ Pur, Dr. Maisch, Ammerbuch, Germany). Peptides were eluted using a gradient from 3.8 to 38% solvent B in solvent A over 120 min at 266 nL per minute flow rate. Solvent A was 0.1% formic acid and solvent B was 79.9% acetonitrile, 20% H_2_O, 0.1% formic acid. Nanoelectrospray was generated by applying 3.5 kV. A cycle of one full Fourier transformation scan mass spectrum (3001750 m/z, resolution of 60,000 at m/z 200, automatic gain control (AGC) target 1 × 106) was followed by 12 data-dependent MS/MS scans (resolution of 30,000, AGC target 5 × 105) with a normalized collision energy of 25 eV. To avoid repeated sequencing of the same peptides, a dynamic exclusion window of 30 sec was used. Raw MS data were processed with MaxQuant software (v1.6.0.1) and searched against the human proteome database UniProtKB with 21,074 entries, released in December 2018. Parameters of MaxQuant database searching were a false discovery rate (FDR) of 0.01 for proteins and peptides, a minimum peptide length of seven amino acids, a first search mass tolerance for peptides of 20 ppm and a main search tolerance of 4.5 ppm. A maximum of two missed cleavages was allowed for the tryptic digest. Cysteine carbamidomethylation was set as a fixed modification, while N-terminal acetylation and methionine oxidation were set as variable modifications. The MaxQuant processed output files can be found in Supplementary Table 2, showing peptide and protein identification, accession numbers, % sequence coverage of the protein, and q-values.

### Pull-downs and immunoblotting

Streptavidin-pulldowns (Fig. 1B) were carried out using stable-cell lines expression SRSF1-12 proteins. Briefly, for each cell line, ∼1 million cells (one well of a 6-well dish, ∼90% confluent) were induced with 0.1µg/mL doxycycline (final) for ∼16hrs, solubilised with 500µL of 1xNLB (1X PBS, 0.3M NaCl, 1% Triton™ X-100, 0.1% TWEEN® 20) + 1x PhosSTOP, sonicated with Bioruptor (30s ON/OFF, 5 cycles on LO) and centrifuged for 10 minutes at ∼20.000 rcf at 4C to remove cellular debris. Biotinylated target proteins were purified with 25µL (slurry) of MyONE-C1 streptavidin beads (Thermo Fisher Scientific, 65002), pre-washed with 1x NLB + 1x PhosSTOP, for 2hrs in the cold-room with end-to-end rotation. Beads were washed 3 times with 500µL of 1x NLB (5 minutes each), bound proteins were eluted with 50µL of 1xLDS sample buffer (Thermo Fisher Scientific, NP0007) + 100mM beta-mercaptoethanol at 95C for 5 minutes. Eluates were loaded on a 4-12% Bis-Tris gel (Thermo Fisher Scientific, NP0322PK2) and transferred to a 0.45µm PVDF membrane (Merck Millipore, IPVH00010) with 10mM CAPS (pH 11) + 10% MeOH, for 900 minutes at 20V. Primary antibodies were used at a dilution of 1:1000 in SuperBlock™ (Thermo Fisher Scientific, 37515). Membranes were incubated with the diluted primaries overnight in the cold-room. SC35 and IgG immunoprecipitations (Fig. 1C) were carried out using a whole-cell extract prepared from wild-type HEK293 cells. Briefly, ∼10 million cells were resuspended with 600µL of 1x NLB + 1x cOmplete™ Protease Inhibitor Cocktail + 1x PhosSTOP, and kept on ice for 15 minutes. The lysate was cleared by centrifugation at ∼20.000 rcf for 10 minutes at 4°C. Clarified lysate was split into two tubes; to one tube 25µL of control IgG1 (Santa Cruz, sc-3877) was added, to the other 2.5µL of SC35 mAb (Sigma-Aldrich, S4045), immune-complexes are allowed to form for 3 hrs in the cold-room with end-to-end rotation. 40µL of Protein G Dynabeads™ (Thermo Fisher Scientific, 10003D, washed and resuspended with 200µL of 1x NLB + PI + PS) was used to pull-down target proteins. Beads were washed three times with 1xNLB, briefly with HSB (50mM Tris.Cl pH 7.4, 1M NaCl, 1% IGEPAL® CA-630, 0.1% SDS, 1mM EDTA) and finally with NDB (50mM Tris.Cl pH 7.4, 0.1M NaCl, 0.1% TWEEN® 20). Bound proteins were eluted with 50µL of 1xLDS sample buffer (Thermo Fisher Scientific, NP0007) + 100mM beta-mercaptoethanol at 80°C for 10 minutes. Immunoblotting was carried out as described for streptavidin pull-downs, except transfer was carried out with 25 mM Tris, 192 mM glycine, 20% (v/v) for 90 minutes at 90V in the cold-room.

Pull-down of truncated SRRM2 proteins (Fig. 2C-E) were carried out using whole-cell lysate prepared from respective HAP1 cell lines. The protocol is essentially identical to SC35 and IgG immunoprecipitations described above, with these notable differences: 1) For pull-downs, 25µL (slurry) of GFP-trap agarose beads were used (Chromotek, gta), incubations were carried out overnight in the cold-room 2) For Fig. 2C and Fig. 2D, 3-8% Tris-Acetate gels (Thermo Fisher Scientific, EA0375PK2) were used, for Fig. 2E a 4-12% Bis-Tris gel was used 3) Gels were run at 80V for 3hrs 4) Transfers were carried out with 10mM CAPS (pH 11) + 10% MeOH for 900 minutes at 20V.

#### Antibodies

ADAR1(Santa Cruz Biotechnology, sc-73408), alpha-Tubulin (Santa CruzBiotechnology, sc-32293), COIL (Cell Signaling Technology, D2L3J, 14168), DHX9 (Abcam, ab183731), FLAG-M2 (Sigma, F3165), GFP (Chromotek, PAGB1), Myc-tag (Cell Signaling Technology, 9B11, 2276), Phospho-Rpb1 CTD (Ser2) (Cell Signaling Technology, E1Z3G, 13499), PNN (Abcam, ab244250), RBM25 (Sigma, HPA070713-100UL), SC-35 (Sigma, S4045), SRSF1 (SF2/ASF, Santa Cruz Biotechnology, sc-33652), SON (polyclonal rabbit, Sigma, HPA023535), SRRM1 (Abcam, ab221061), SRRM2 (Thermo Fisher Scientific, PA5-66827), SRSF2 (Thermo Fisher Scientific PA5-92037), SRSF7 (MBL, RN079PW), U1-70K (Santa Cruz Biotechnology, sc-390899), U2AF65 (Santa Cruz Biotechnology, sc-53942)

##### Note on SC35/SRSF2 antibodies

There are many commercially available antibodies that are labeled as ‘SC35’, however only some of them are actually clones of the original SC-35 antibody reported by Fu and Maniatis in 1990. These are: s4045 from Sigma-Aldrich, sc-53518 from Santa Cruz Biotechnology and ab11826 from Abcam. Some antibodies are sold as ‘SC35’ antibodies, but they are antibodies specifically raised against SRSF2. These are: ab204916 and ab28428 from Abcam and 04-1550 from Merck (can be found with the clone number 1SC-4F11). Neither list is exhaustive.

### Phylogenetic analysis

Unless indicated otherwise, all data analysis tasks were performed using Python 3.7 with scientific libraries Biopython (75), pandas (76), NumPy (77), matplotlib (78) and seaborn. Code in the form of Jupyter Notebooks is available in GitHub repository: https://github.molgen.mpg.de/malszycki/SON_SRRM2_speckles Vertebrate SRRM2, SON, PRPF8, SRRM1, RBM25, Pinin and Coilin orthologous protein datasets were downloaded from NCBI’s orthologs and supplemented with orthologues predicted for invertebrate species. For this purpose, OrthoFinder (79) was used on a set of Uniprot Reference Proteomes. Invertebrate orthologues were then mapped to NCBI RefSeq to remove fragmentary and redundant sequences. The resulting dataset was manually curated to remove evident artefacts lacking conserved domains or displaying striking differences from closely related sequences. Protein lengths were plotted using the seaborn package and descriptive statistics calculated using the pandas package. In order to resolve phylogenetic relationships between species contained in SRRM2 and SON datasets, organism names were mapped to the TimeTree (80) database. Disorder probability was predicted using IUPred2A (81) and MobiDB-Lite (82) and plotted as a heatmap using matplotlib.

---

**Fig. S1.**
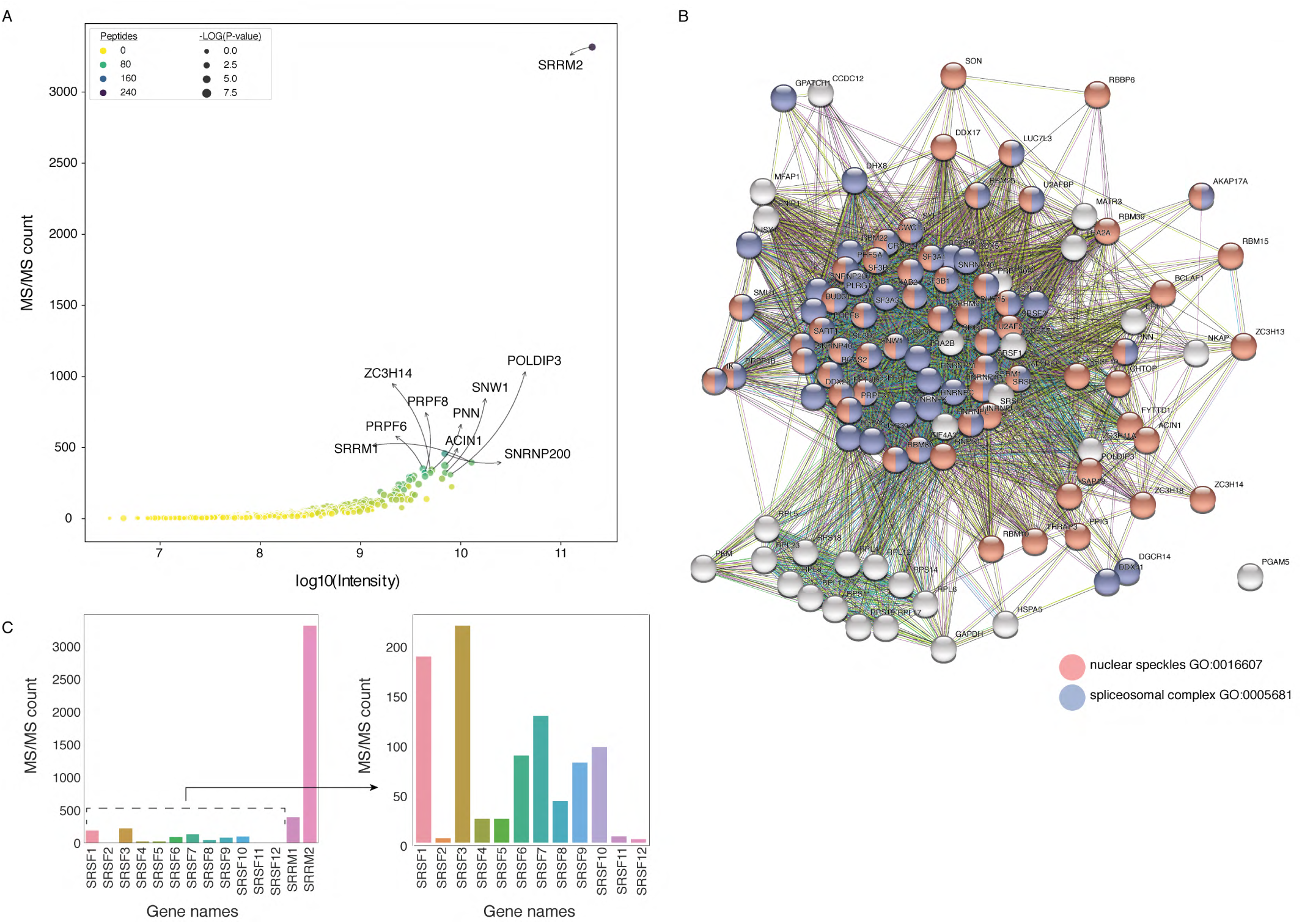
SC35 pull-down followed by MS identifies SRRM2 as the top hit and multiple spliceosomal components are co-purified together with SRRM2. **(A)** The log of intensities is plotted against detected MS/MS spectra for significantly enriched proteins in the SC35 immunoprecipitations. SRRM2 together with nine most enriched proteins are annotated on the plot. **(B)** SC35 targets in the top quartile (108 proteins) are submitted into the STRING database and a network representation is shown. Red circles depict nuclear speckle GO-term category and blue circles depict spliceosomal complex GO-term category. (C) The MS/MS spectral counts for SRSF1 to 12 are plotted together with SRRM1 and SRRM2 (left) and without SRRM1 or SRRM2 (right) show a significant enrichment for SRRM2 amongst other SR-proteins.

**Fig. S2.**
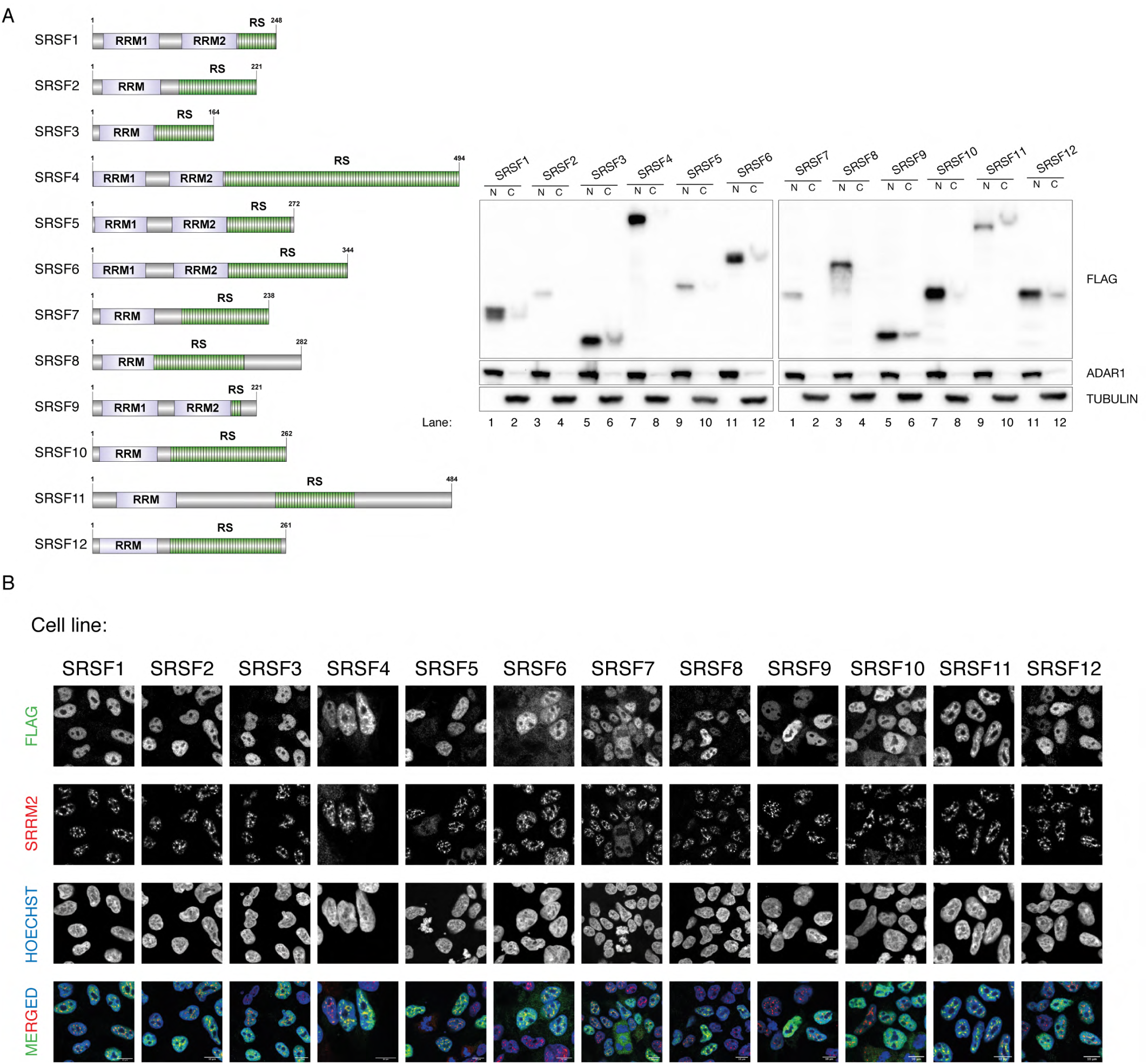
A validation for all stable cell lines with transgenic SRSF proteins show only weak localization to the NS. **(A)**The domain and size representation of SRSF proteins are shown (left). The inducible stable cell lines are characterized using PAGE (right). The nuclear (N) and cytoplasmic (C) fractions are used to show the major localization to the nucleus is unaffected when the SRSF proteins are tagged and ectopically expressed. ADAR1 is used as a nuclear marker whereas TUBULIN is used as a cytoplasmic marker **(B)**The Immunofluorescence experiment is performed by using FLAG-M2 for the tagged SRSF protein and by using SRRM2 (pRb) antibody as an NS marker. Overall NS localisation of SRSF proteins is noted, but in a more diffused pattern in comparison to SRRM2. Scale bars = 10µm.

**Fig. S3.**
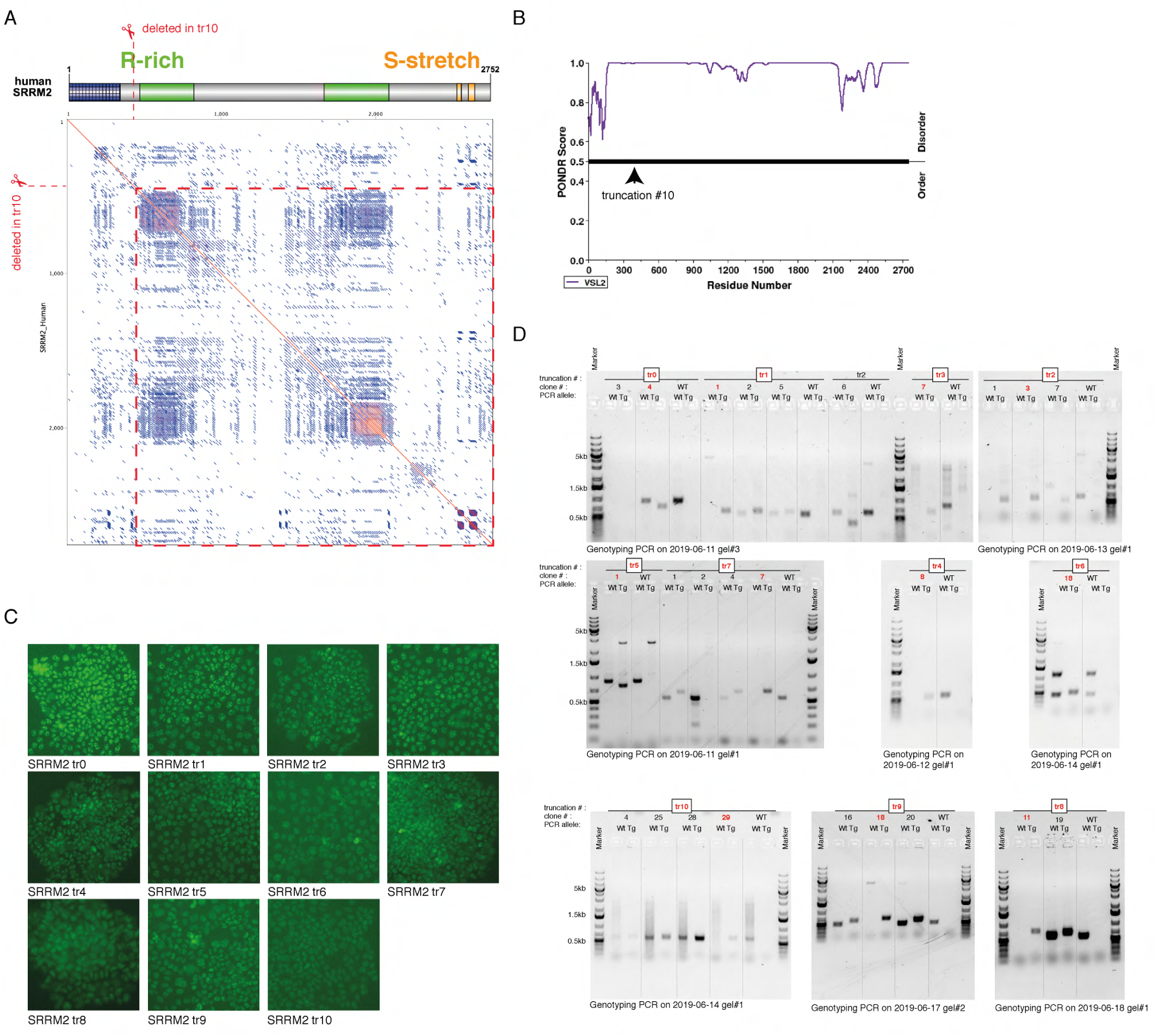
The strategy for making the truncating mutations of SRRM2. **A)**The position of the deepest truncation (tr10) is shown with respect to annotated domains of the protein as well as the dot matrix of the SRRM2 protein sequence. Repetitive regions appear as densed blue squares on the matrix. **(B)**The PONDR score for disordered regions of SRRM2 protein is displayed with the arrow head pointing to the position of truncation 10.**(C)** A summary of GFP fluorescence images for truncating mutations before the colony picking is shown. **(D)** The results of genotyping PCRs are summarized (Wt: PCR carried out using the oligos for wild type allele; Tg:PCR carried out using the oligos for wild type allele. Genotyping oligos and the expected PCR product sizes are listed in Supplementary Table 1). Every clone used in this study is highlighted with red text color.

**Fig. S4.**
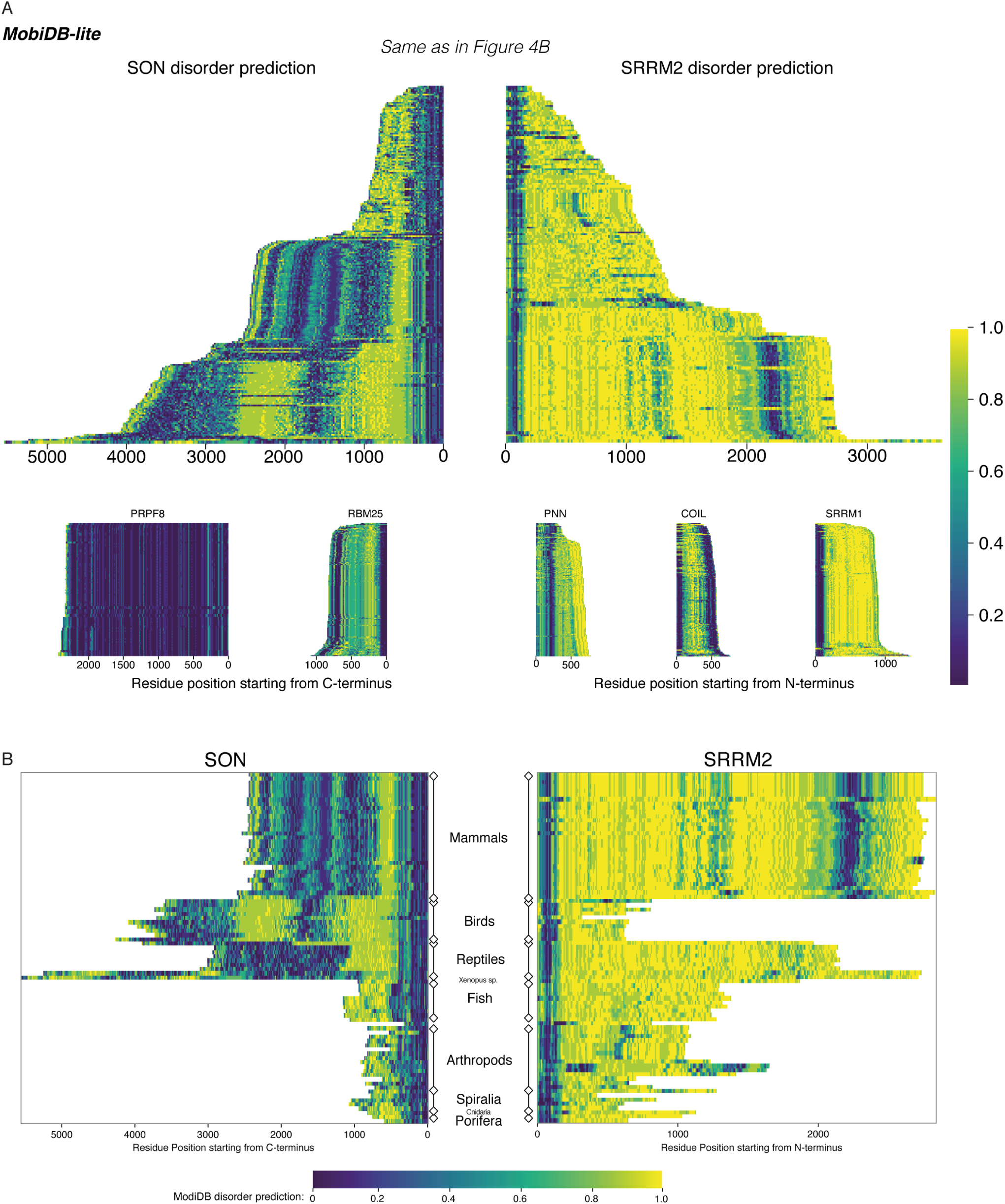
SON and SRRM2 are rapidly evolving and largely disordered proteins. **(A)**The disorder probability of SON and SRRM2 is shown the same way as in Fig. 4B calculated using MobiDB-Lite (up). The same analysis is also carried out on a spliceosomal core protein PRPF8, on other NS-associated proteins RBM25, PNN and SRRM1, and on a different nuclear body (Cajal bodies) scaffold protein COIL. The length for the highly disordered protein SRRM1 does not vary to a similar extent as SON or SRRM2, indicating the length changes are not a direct consequence of disorderedness. **(B)**The disorder probability of SON and SRRM2 is shown in phlogenetically matched order using MobiDB-Lite for disorder prediction. In mammals the length of both proteins seem to be fixed, whereas birds have shorter SRRM2 compared to mammals but longer SON. Strikingly, Xenopus tropicalis and laevis have both SON and SRRM2 increased in length.

**Fig. S5.**
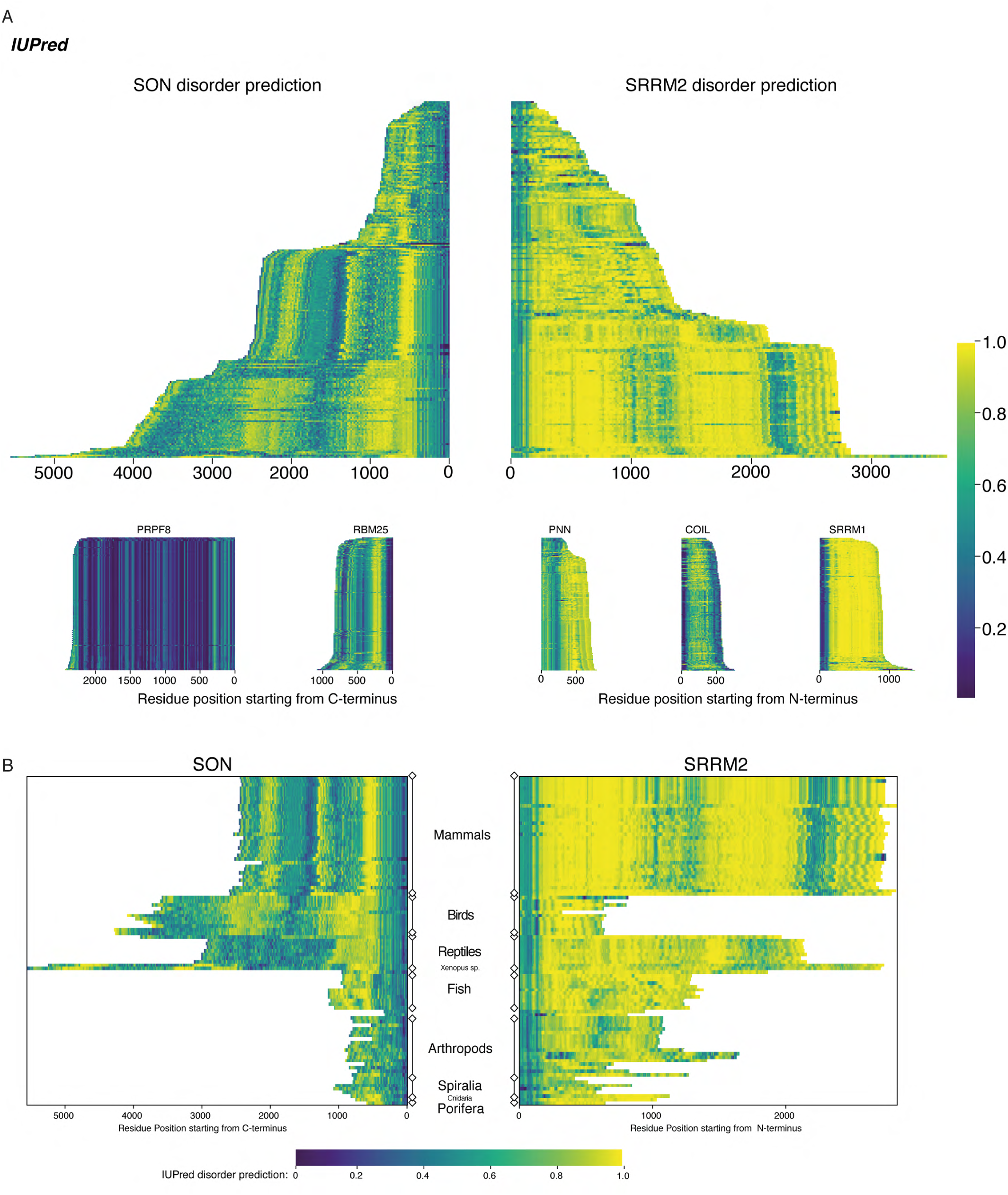
SON and SRRM2 are rapidly evolving and largely disordered proteins. **(A)**The disorder probability of SON and SRRM2 is shown similar to Fig. 4B and Supplementary Fig. 4A calculated using IUPred2A. Both disorder prediction tools resulted in similar plots. **(B)**The disorder probability of SON and SRRM2 is shown in phlogenetically matched order using IUPred2A for disorder prediction, similar to Supplementary Fig. 4B. Both disorder prediction tools resulted in similar plots.

**Fig. S6.**
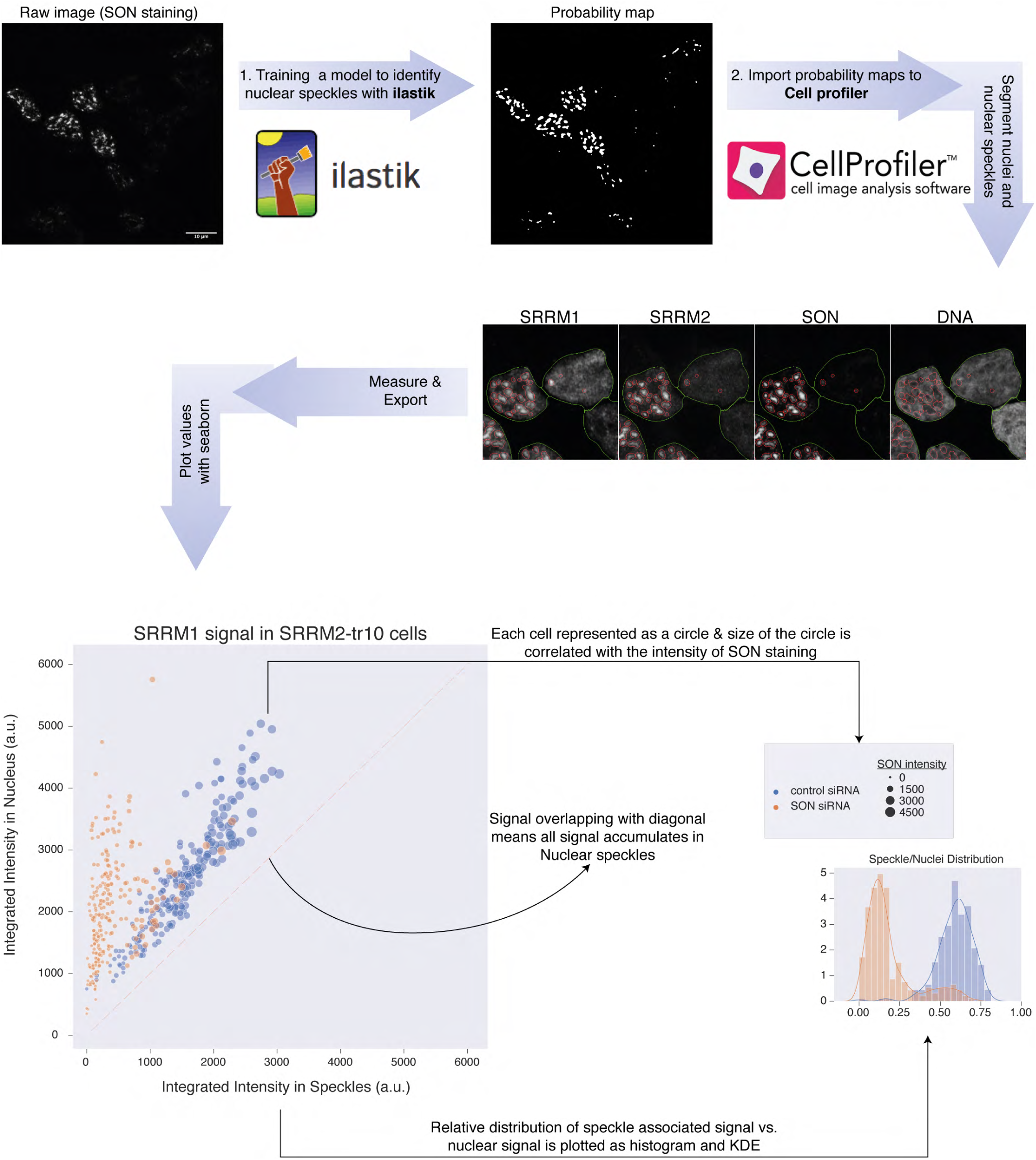
Training a machine learning method for detection of NS. The pipeline used for the quantification of protein localisation in NS under various conditions is depicted here. The probability maps generated with ilastik are imported into CellProfiler analysis software for segmentation and analysis. The numerical values obtained from CellProfiler are then used for the plots which are shown in Fig. 5, Supplementary Fig. 8 and Supplementary Fig. 9. The green line marks the nuclear boundaries and the red circles are detected as NS by the algorithm.

**Fig. S7.**
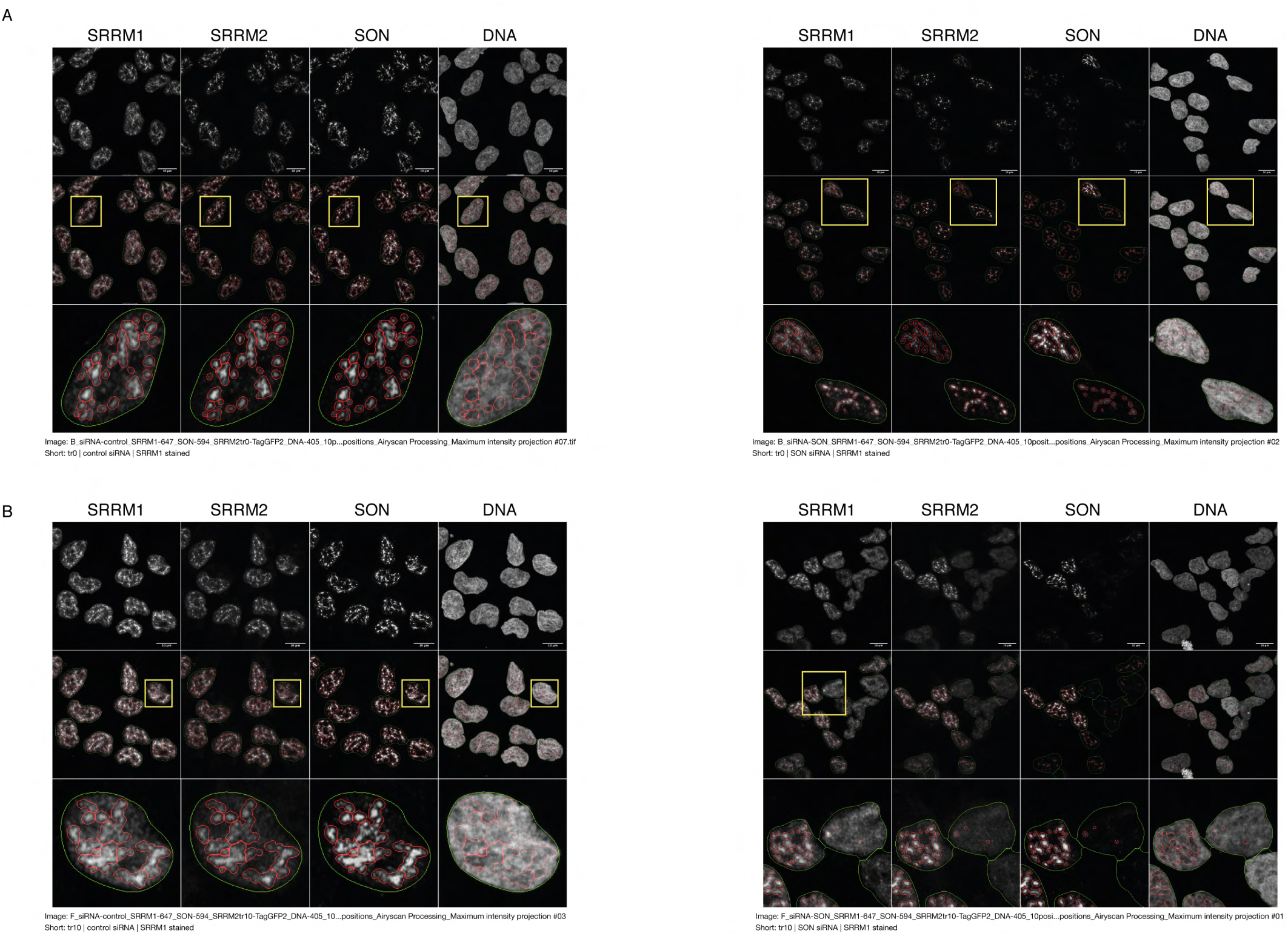
Examples of the trained module for detection of NS on different antibody stainings indicate the model predicts NS robustly for each stained protein. **(A)** The outcome of the trained model on recognition of NS is shown for SRRM2^tr0^ HAP1 cells treated with control siRNA (left) or SON siRNA (right). **(B)**The outcome of the trained model on recognition of NS is shown for SRRM2^tr10^ HAP1 cells treated with control siRNA (left) or SON siRNA (right). The green line marks the nuclear boundaries and the red circles are detected as NS by the algorithm. Scale bars = 10µm

**Fig. S8.**
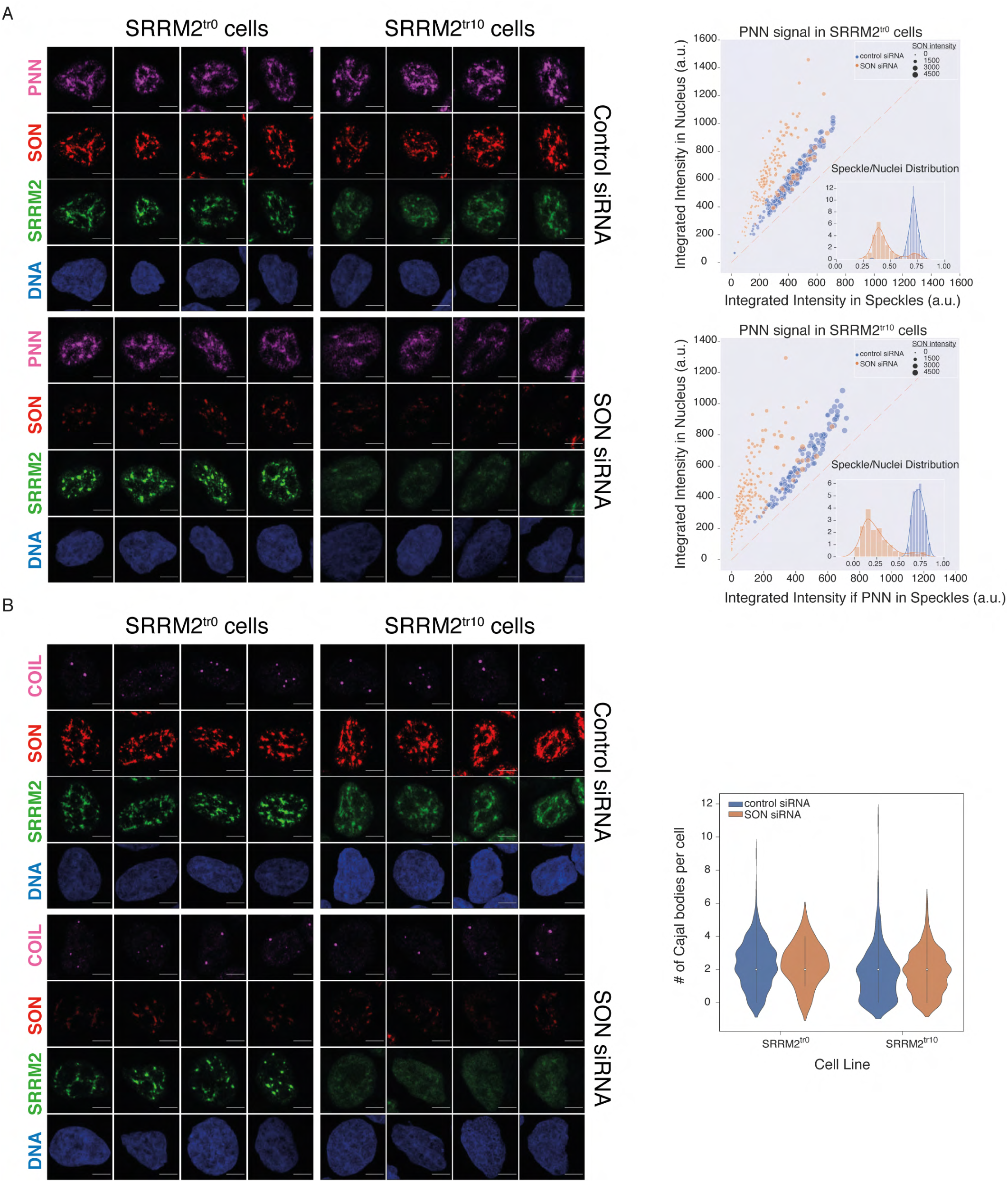
Depletion of SON in SRRM2tr10 cells leads to loss of NS but not of Cajal bodies. **(A)** PNN IF signal is shown for four individual cells in each siRNA treatment (control or SON siRNA) in SRRM2^tr0^ and SRRM2^tr10^ HAP1 cells. The NS localization of PNN is lost upon SON knock-down in SRRM2tr10 cells. The quantification of the PNN signal within the nucleus is plotted against the PNN signal within NS (right panel) using ilastik to train detection of NS and CellProfiler for quantification on 10 imaged fields with 63X objective (in SRRM2tr0 cells control n=346, SON-KD n=172; in SRRM2tr10 cells control n=138, SON-KD n=199). Each circle represents a cell and the size of the circles is proportionate to the signal intensity of SON. Inset shows the distribution of the ratio of signal detected in NS over signal detected in the nucleus of each cell. **(B)** COIL IF signal is shown for four individual cells in each siRNA treatment (control or SON siRNA) in SRRM2^tr0^ and SRRM2^tr10^ HAP1 cells. There is no significant change in the localization or of the signal intensity of COIL upon SON knock-down. The quantification of the number of Cajal bodies (based on COIL signal) within the nucleus is shown in the right panel (in SRRM2^tr0^ cells control n= 337, SON-KD n=63; in SRRM2^tr10^ cells control n=309, SON-KD n=244). Scale bars = 5µm

**Fig. S9.**
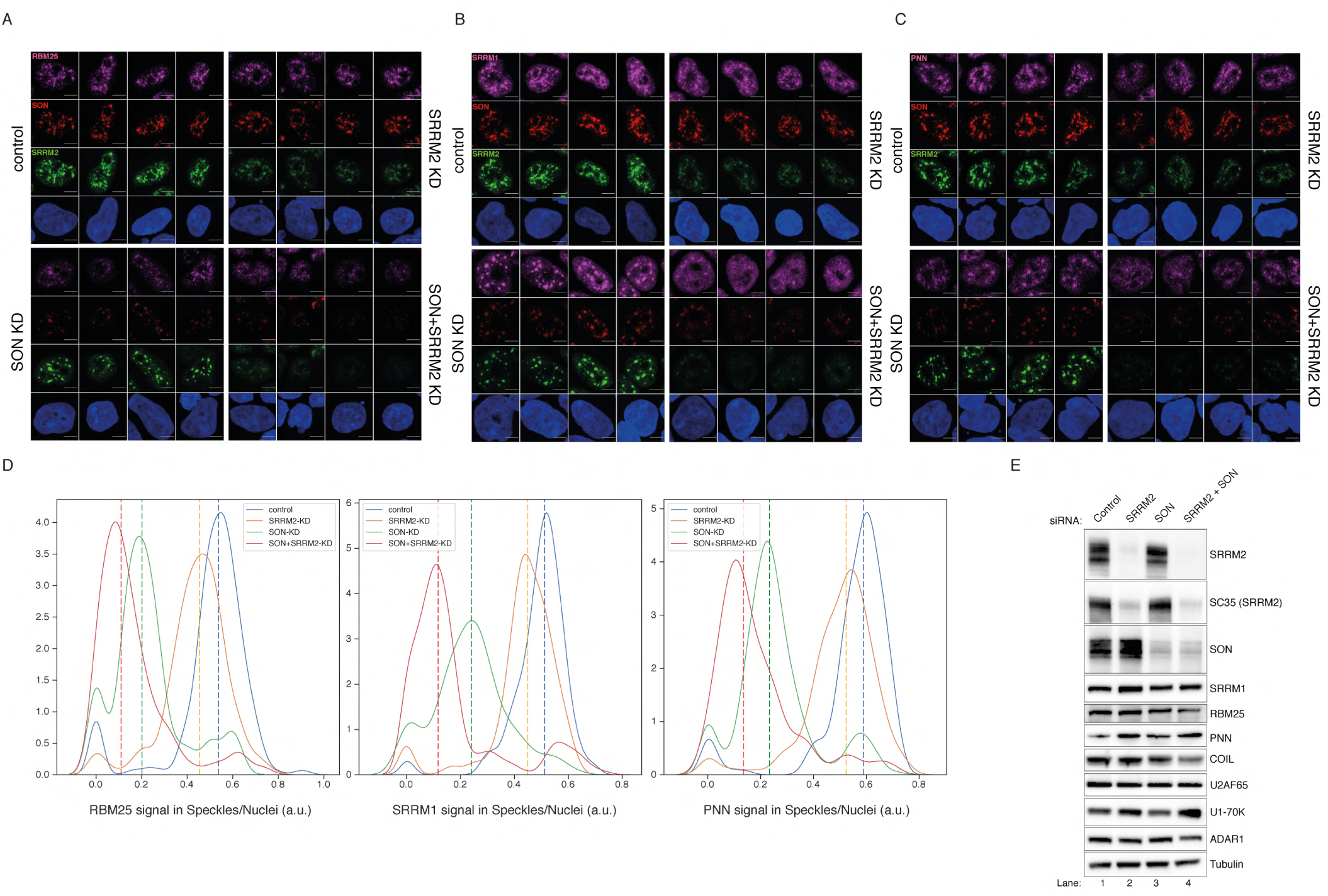
Co-depletion of SON and SRRM2 in SRRM2^tr0^+GFP HEK293 cells leads to loss of NS. **(A)** RBM25 IF signal is shown for four individual cells in each siRNA treatment (control, SRRM2, SON or SRRM2 SON siRNA) in SRRM2^tr0^+GFP HEK293 cells. RBM25 signal diffuses out of NS upon SON depletion but is completely lost only in the double knock-down cells. **(B)** SRRM1 IF signal is shown for four individual cells in each siRNA treatment (control, SRRM2, SON or SRRM2 SON siRNA) in SRRM2^tr0^+GFP HEK293 cells. SRRM1 signal diffuses out of NS upon SRRM2 depletion and is mostly diffused in the nucleus in the double knock-down cells. **(C)**PNN IF signal is shown for four individual cells in each siRNA treatment (control, SRRM2, SON or SRRM2 SON siRNA) in SRRM2tr0+GFP HEK293 cells. PNN signal diffuses out of NS upon SON depletion and is mostly diffused in the nucleus in the double knock-down cells. Scale bars = 5µm **(D)** Distribution plots showing the ratio of signal detected in NS over signal detected in the nucleus of each cell, in each condition. The dashed line indicates the median ratio in each condition, similar to Fig. 5C (for RBM25 staining in control siRNA n=321, in SON-KD n=187; in SRRM2-KD n=249, in double-KD n=202; for SRRM1 staining in control siRNA n=155, in SON-KD n=150; in SRRM2-KD n=197, in double-KD n=262; for PNN staining in control siRNA n=204, in SON-KD n=264; in SRRM2-KD n=299, in double-KD n=229). The double knock-down of SRRM2 and SON leads to the most significant loss of signal localised to the NS for RBM25, SRRM1 and PNN. **(E)**he diffusion of RBM25, SRRM1 or PNN out of the NS is not caused by the down-regulation of the protein levels as shown by western blot (see Lanes 1 and 4).

**Fig. S10.**
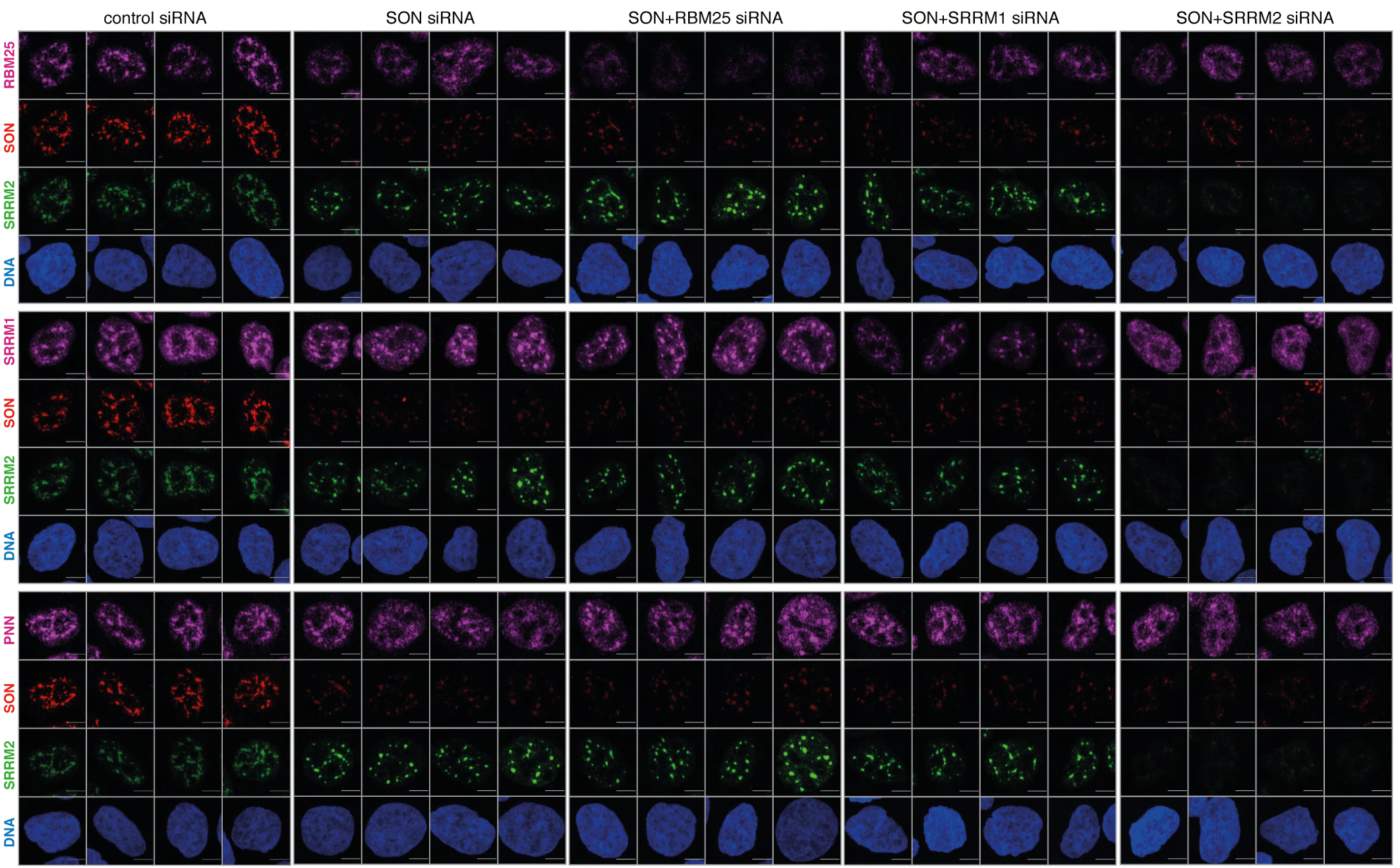
Co-depletion of SON with RBM25 or SRRM1 in SRRM2^tr0^ HAP1 cells does not lead to loss of spherical NS. Knock-down experiments are performed similar to Supplementary Fig. 9, but in SRRM2^tr0^ HAP1 cells, however this time SON is also co-depleted together with RBM25 or SRRM1. Depletion of SON together with RBM25 or SRRM1 does not lead to loss of NS as can be seen in GFP signal of SRRM2, indicating the depletion of any NS-associated protein together with SON does not lead to dissolution of NS. The NS are lost only when SRRM2 and SON are co-depleted. Scale bars = 5µm.

