## Supplementary figures and images for "SON and SRRM2 form nuclear speckles in human cells"

### Summary Figure

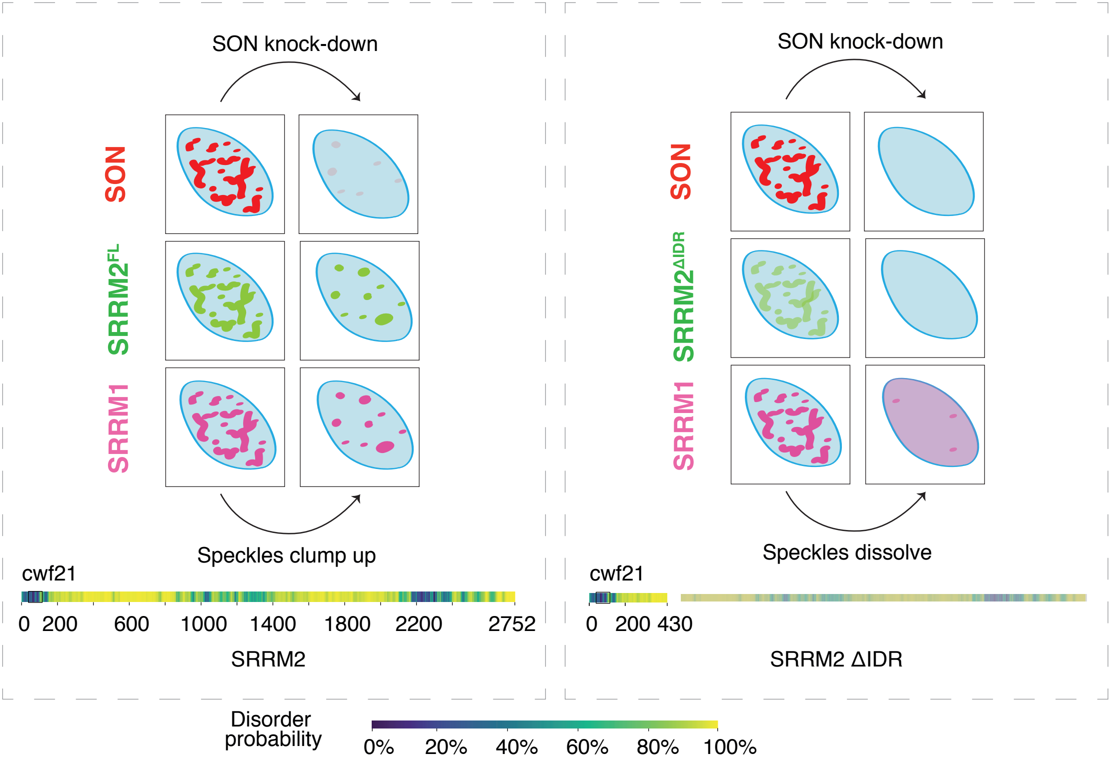
